# Pattern of Circulating Mesenchymal Stromal Cells and Hematopoietic Progenitor and Stem Cells in the Peripheral Blood of Trauma Patients with and without Hemorrhagic Shock

**DOI:** 10.64898/2026.03.28.714706

**Authors:** V Priya Dharshani, Sanjeev Kumar Bhoi, Subhradip Karmakar, Tej Prakash Sinha

**Affiliations:** JPN APEX TRAUMA CENTRE ALL INDIA INSTITUTE OF MEDICAL SCIENCES, NEW DELHI, INDIA; ALL INDIA INSTITUTE OF MEDICAL SCIENCES, NEW DELHI, INDIA

**Keywords:** hemorrhagic shock, trauma, hematopoietic stem cells, mesenchymal stromal cells, progenitor cell mobilisation, cytokines, inflammation, flow cytometry, bone marrow response, critical care

## Abstract

Circulating stem and progenitor cells (SPCs), including mesenchymal stromal cells (MSCs) and hematopoietic stem/progenitor cells (HSPCs), are mobilised after tissue injury but their temporal behaviour after hemorrhagic shock (HS) and relationship to cytokine milieus and outcome remain unclear. In a prospective observational cohort at JPN Apex Trauma Centre, AIIMS, New Delhi we studied 100 participants: 50 trauma patients with hemorrhagic shock and traumatic brain injury (HS index group), 25 trauma patients without HS, and 25 minor-injury controls. Peripheral blood was collected at admission (day 0) for all groups and additionally at days 3, 7 and 14 for the HS group. PBMCs were phenotyped by flow cytometry (HSPC markers: CD45, CD123, CD38, CD34; MSC markers: CD105, CD73, CD90) and serum SDF-1, VEGF-A, EGF, GRO-α and GRO-β, GM-CSF and G-CSF were measured by ELISA; group and time effects were evaluated with mixed-effects models and correlations by Spearman tests (two-tailed p<0.05). At admission, trauma patients without HS had significantly higher MSC and HSPC-like populations versus controls (p<0.0001). In the HS cohort SPC percentages rose modestly at day 0-3 then declined sharply by days 7-14 (time effect p<0.0001); non-survivors exhibited significantly higher early SPC and cytokine levels that persisted until death while survivors showed an early rise followed by decline (outcome and time interaction p<0.0001). All cytokines were up-regulated in trauma groups, peaked at day 0-3 in HS patients, and correlated positively with SPC counts (notably SDF-1, VEGF-A, G-CSF, Gro-α and GM-CSF; Spearman p<0.05); higher early SPC and cytokine signatures associated with greater organ dysfunction (higher SOFA) and with timing of sepsis. These findings indicate that trauma provokes an early SPC and cytokine response that in HS is followed by later decline, and that persistent early elevation predicts worse outcomes, suggesting serial SPC and cytokine profiling may have prognostic value and identify an early therapeutic window for regenerative or immunomodulatory interventions.

## Introduction

Trauma and hemorrhagic shock (HS) are major causes of morbidity and mortality worldwide; early mortality after HS remains high, frequently occurring within the first 24 hours. Elevated systemic inflammation and sympathetic activation after severe injury are linked to hematopoietic dysfunction, anemia, and increased infection risk (Kumar & Bhoi, 2016; Kumar et al., 2019). Hematopoietic stem and progenitor cells (HSPCs) resident in bone marrow and mesenchymal stromal cells (MSCs) comprise circulating SPC populations that are intermittently mobilised into peripheral blood following tissue injury (Thomas et al., 2002). MSCs exert both regenerative and immunomodulatory effects and MSC-derived extracellular vesicles reduce vascular leak and inflammation in preclinical models of HS (Pati et al., 2011; Potter et al., 2018). HSPCs (including common myeloid progenitors and granulocyte-macrophage progenitors) can contribute pro-angiogenic activity that may assist tissue repair (Cook et al., 2013; Suárez-Monteagudo et al., 2009).

Prior clinical and preclinical reports describe early increases of circulating SPCs within hours after injury with declines by 24–48 h (Kumar et al., 2016), but sample sizes were small and synchronised profiling with multiple cytokines and clinical outcomes over two weeks has been limited. Moreover, how SPC mobilization trajectories differ between trauma patients with and without HS and whether specific cytokine milieus (e.g., SDF-1α, VEGF, EGF, GRO-α/β, GM-CSF, G-CSF) track with mobilization or outcome has not been fully clarified. Understanding these dynamics could identify prognostic biomarkers and therapeutic windows for SPC-based or cytokine-directed therapies.

There is limited, systematically collected data describing synchronised, serial SPC mobilization (MSCs and HSPCs) in trauma patients with HS compared to trauma without HS and controls, linked to serial cytokine profiling and clinical outcomes.

The core objectives of this study is to evaluate temporal patterns (days 0, 3, 7, 14) of circulating MSCs and HSPCs and associated cytokine profiles in three groups: trauma patients with hemorrhagic shock, trauma patients without HS, and minor-injury controls, and to correlate these patterns with clinical outcome (survival vs. death) and organ dysfunction and additionally with the serum cytokine (SDF-1, VEGF-A, EGF, Gro-α, Gro-β, GM-CSF, G-CSF) levels.

We hypothesised that severe trauma induces an early rise in circulating mesenchymal stromal cells (MSCs) and hematopoietic stem and progenitor cells (HSPCs); their levels along with key cytokines such as SDF-1, VEGF-A, EGF, Gro-α, Gro-β, GM-CSF, G-CSF, decline over time post injury and higher early levels of these cells and cytokines are associated with poor clinical outcomes.

## Methods

1. Study Design and Setting: Prospective observational cohort study conducted over three years at the Department of Emergency Medicine (sample collection) and Department of Biochemistry (bench work), All India Institute of Medical Sciences (AIIMS), New Delhi. Ethical clearance was obtained from the institutional ethics committee; written informed consent was obtained from all participants or legally authorised representatives.
2. Participants: Sample Size and Groups- Total n=100.

Index group: Trauma patients with hemorrhagic shock and traumatic brain injury (TBI), n=50.

Comparator group A: Trauma patients without hemorrhagic shock (no TBI), n=25. Comparator group B: Minor-injury patients, n=25.

Inclusion criteria: age 18–45 years; admission within 8 hours of injury; systolic blood pressure ≤90 mm Hg for HS group; written informed consent.

Exclusion criteria: <18 or >45 years; admission >8 hours after injury; pre-existing significant illness (chronic organ disease, immunosuppression, malignancy, etc.).

Clinical data recorded included demographics, dates of admission/discharge, routine clinical variables, 15-day in-hospital mortality, and organ dysfunction quantified by the SOFA score (respiratory, cardiovascular, hepatic, coagulation, renal, neurological subscores).

3. Sample Collection and Processing:

At each time point (day 0 on admission, day 3, day 7, day 14), 4–5 mL peripheral venous blood was collected into appropriate anticoagulant tubes. Plasma/serum were separated and aliquoted for cytokine assays; PBMCs were isolated for flow cytometric phenotyping.

PBMC Isolation: Whole blood samples were incubated at room temperature for 30 min and centrifuged at 1600 × g for 10 min, after which the supernatant was discarded. Residual red blood cells were lysed using 1X RBC lysis buffer for 10–15 min at room temperature, followed by centrifugation at 10,000 × g for 20 min. The resulting pellet was transferred to a 1.5 mL tube, subjected to a second lysis for 10 min, and centrifuged at 2,500 × g for 3 min. The pellet was washed twice with 1X PBS (2,500 × g, 3 min each) to obtain peripheral blood mononuclear cells (PBMCs). PBMCs were counted using a hemocytometer, and 1 × 10⁶ cells were used per staining. Cells were fixed in 2% paraformaldehyde and stored at 4 °C; before acquisition, fixed samples were incubated for 10 min at room temperature and washed twice in PBS.

4. Flow cytometry- Identification and Quantification of SPCs:

Flow cytometric analysis was performed using a BD LSRFortessa™ Cell Analyser, and data were analyzed with FlowJo software. The following fluorochrome-conjugated mouse anti-human antibodies were used for surface staining: CD38 (BV711, Cat. No. 563965), CD105 (PerCP-Cy5.5, Cat. No. 560819), CD123 (BUV395, Cat. No. 567277), CD73 (APC, Cat. No. 560847), CD14 (FITC, Cat No. 563440), CD34 (PE, Cat. No. 563119), CD45 (BV510, Cat. No. 563204), CD90 (BV421, Cat. No. 562556), and CD20 (APC-H7, Cat. No. 560853).

Staining Protocol: 100 µL PBMC suspension (∼1 × 10^6^ cells) per tube. Antibodies added per manufacturer-recommended dilutions and incubated on ice in the dark for 45 min. Cells washed twice with BD Horizon Brilliant Stain Buffer, centrifuged (5 min at 1,500 rpm, 4°C). Final resuspension in BD FACS buffer and acquisition on BD LSRFortessa. Fixed samples acquired after re-equilibration as described above.

Laboratory staff performing flow cytometry were blinded to patient clinical outcomes. Instrument settings and compensation were performed using single-stained compensation beads. Fluorescence-minus-one (FMO) controls were used to determine gate thresholds and percentages were reported relative to the parent population. Data were analysed in FlowJo (v10).

Gating: Flow cytometry analysis was performed on mononuclear cells. Viable singlet cells were selected by FSC gating to exclude debris and FSC-A vs FSC-H to exclude doublets. HPSCs were identified by flow cytometry as viable singlet, CD45^+^, lineage-negative (CD14^-^, CD20^-^) CD34^+^ cells. Primitive HSC-enriched cells were defined as CD34^+^, CD38^-^ with further enrichment by CD90 expression (CD34^+^ CD38^-^ CD90^+^), while committed progenitors were defined as CD34^+^ CD38^+^. CD123 was included to further phenotype progenitor subsets. MSCs were defined as viable singlet cells negative for hematopoietic markers (CD45^-^, CD14^-^, CD20^-^, CD34^-^) and positive for canonical MSC markers CD105, CD73 and CD90.

5. Cytokine Quantification (ELISA):

Serum total protein quantified by bicinchoninic acid (BCA) assay; 100 µg total protein per sample used for ELISA. Commercial kits (ImmunoTag) were used according to manufacturers’ protocols: SDF-1 (Cat. ITLK01183), VEGF-A (ITLK01129), EGF (ITLK01266), GRO-α (ITLK02078), GRO-β (ITLK02079), GM-CSF (ITLK01028), G-CSF (ITLK01025). Absorbance read at 450 nm. Standard curves constructed and concentrations interpolated. All samples run in duplicate.

6. Statistical Analysis:

Data were analyzed with GraphPad Prism v10. Continuous data were tested for normality using the Shapiro–Wilk test. For normally distributed repeated measures, two-way repeated measures ANOVA (group × time) with post hoc Bonferroni corrections was used; for non-normal repeated data, Friedman test with Dunn’s multiple comparisons was applied. Between-group comparisons at single time points used Student’s t-test or Mann–Whitney U test as appropriate. Correlations between cytokines and SPC counts used Spearman’s rank correlation. Survival comparisons used Kaplan–Meier curves and log-rank tests when appropriate. A two-tailed p < 0.05 was considered statistically significant. Exact p-values, test statistics, and effect sizes are reported in results.

## Results

One hundred patients were enrolled- 25 minor-injury controls, 25 trauma patients without hemorrhagic shock (HS), both comparator group, and 50 trauma patients with HS, index group. Blood from comparator groups (controls and trauma without HS) was collected at admission (day 0) only; index-group samples were collected at day 0 (admission), day 3, day 7 and day 14. Laboratory staff were blinded to clinical outcome.

Circulating SPCs - Trauma without HS versus controls (Comparator Group) on Day 0: At admission (day 0) trauma patients without HS demonstrated a clear mobilization of both mesenchymal stromal cell (MSC-like) and hematopoietic stem/progenitor cell (HSPC-like) populations relative to minor-injury controls (Figure 01, Figure 04) . This early elevation was consistent across the marker panels used (CD105, CD73, CD90 for MSC-like cells; CD123, CD38, CD45 for HSPC-like cells) and was judged statistically significant in group comparisons performed at day 0.

**Figure 01:**
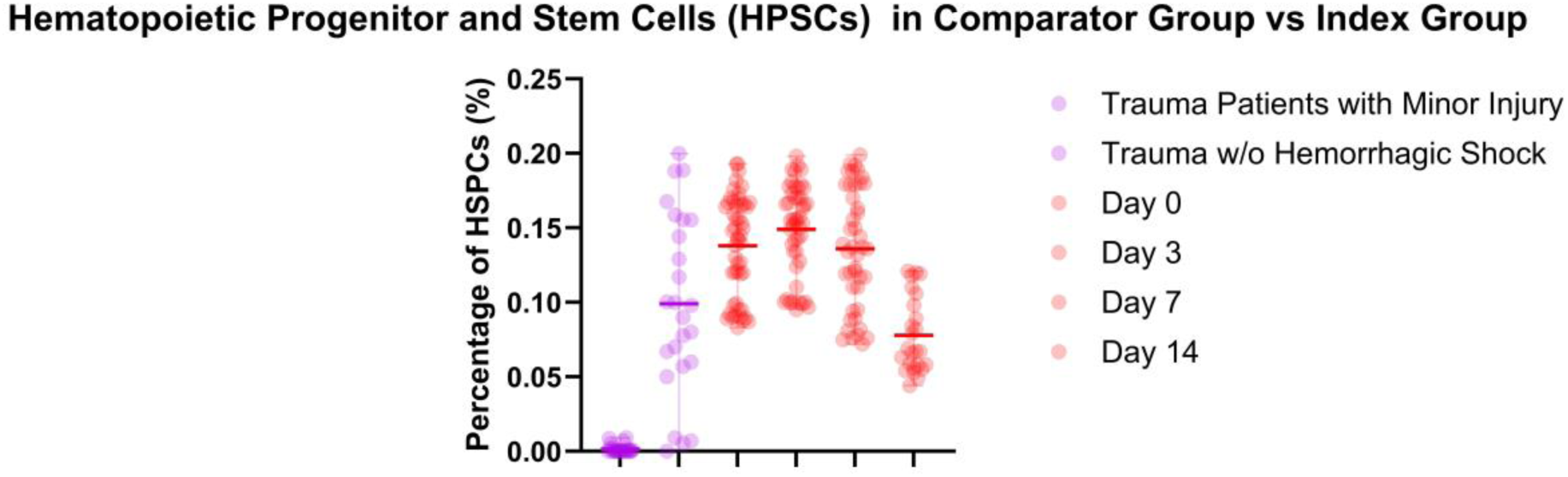
Percentage of Hematopoietic Progenitor Stem Cells (HPSCs) in Peripheral Human Blood. n=25 (Patients with Minor Injury), n=25 (Trauma Patients without Hemorrhagic Shock), n=50 (Trauma Patients with Hemorrhagic Shock and Traumatic Brain Injury); ****p<0.0001.

Longitudinal SPC dynamics in hemorrhagic shock (index group): Among the 50 HS patients the time course of circulating SPCs followed a biphasic pattern (Figure 02, Figure 05):

- Days 0–3: modest but measurable elevations in both MSC- and HSPC-like populations compared with controls.
- Days 7–14: a clear, moderate-to-sharp decline in both MSC and HSPC percentages relative to the day-0/day-3 values.

**Figure 02:**
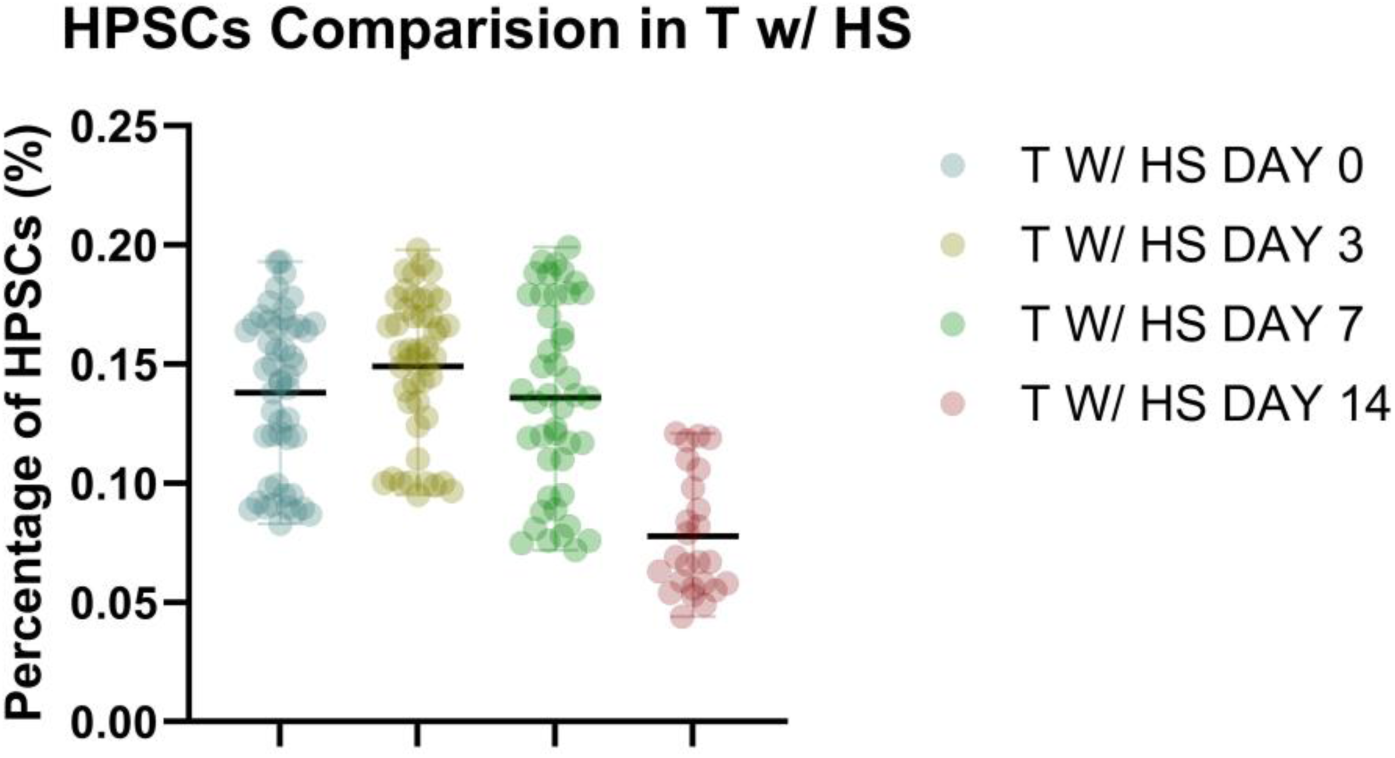
Percentage of Hematopoietic Progenitor Stem Cells (HPSCs) in Peripheral Human Blood of Index Group. n=50 (Trauma Patients with Hemorrhagic Shock and Traumatic Brain Injury); ****p<0.0001.

A mixed-effects analysis indicated a significant time effect (early rise followed by later decline) and a significant group and time interaction compared with the non-HS trauma group (i.e., the temporal trajectory in HS patients differed from the single time-point elevation seen in non-HS trauma patients).

Outcome stratification: survivors versus non-survivors (Index Group): When HS patients were stratified by 15-day outcome, distinct patterns emerged (Figure 03, Figure 06):

- Non-survivors showed significantly higher percentages of circulating MSC- and HSPC-like cells at early time points (day 0 and day 3) and these elevated levels persisted up to the time of death.
- Survivors also showed early (day 0–3) elevations, but these declined by day 7 and day 14.

**Figure 03:**
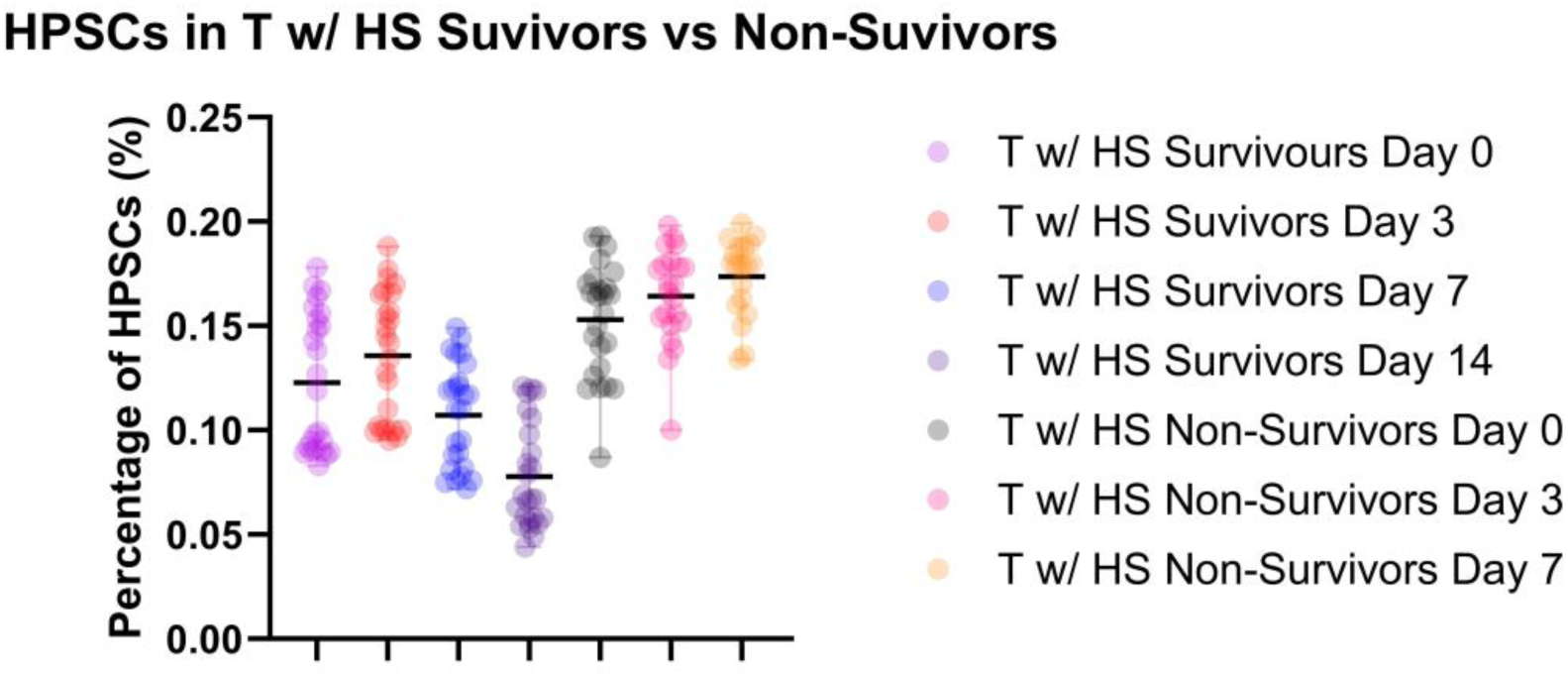
Percentage of Hematopoietic Progenitor Stem Cells (HPSCs) in Peripheral Human Blood among survivors and non-survivors of hemorrhagic shock, n=50 (Trauma Patients with Hemorrhagic Shock and Traumatic Brain Injury); ****p<0.0001.

**Figure 04:**
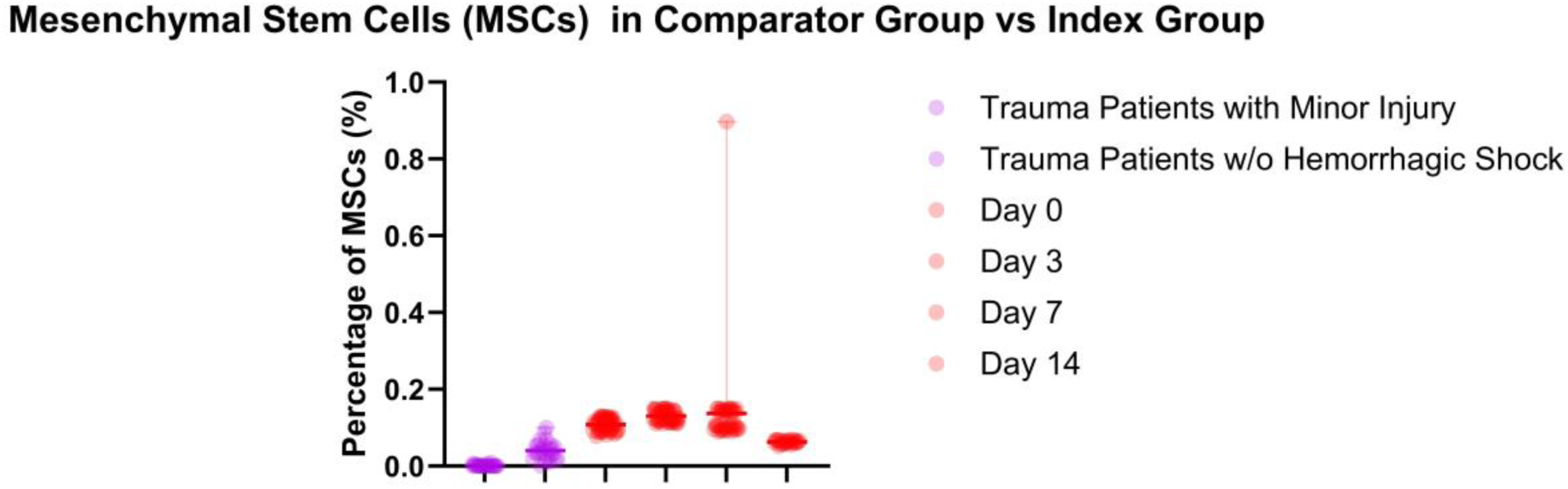
Percentage of Mesenchymal Stromal Cells in Peripheral Human Blood. n=25 (Patients with Minor Injury), n=25 (Trauma Patients without Hemorrhagic Shock), n=50 (Trauma Patients with Hemorrhagic Shock and Traumatic Brain Injury); ****p<0.0001.

**Figure 05:**
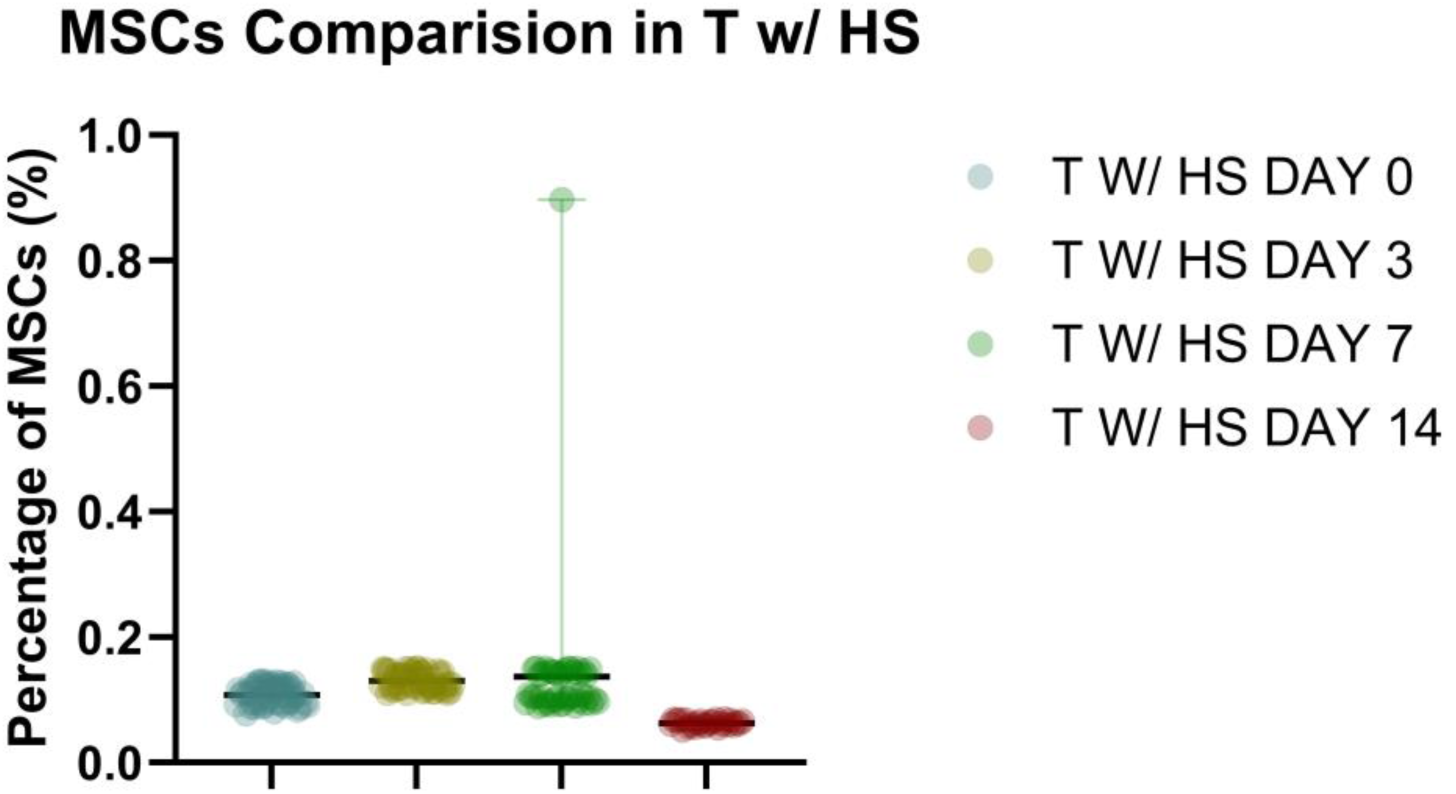
Percentage of Mesenchymal Stromal Cells in Peripheral Human Blood of Index Group. n=50 (Trauma Patients with Hemorrhagic Shock and Traumatic Brain Injury); ***p<0.0007.

**Figure 06:**
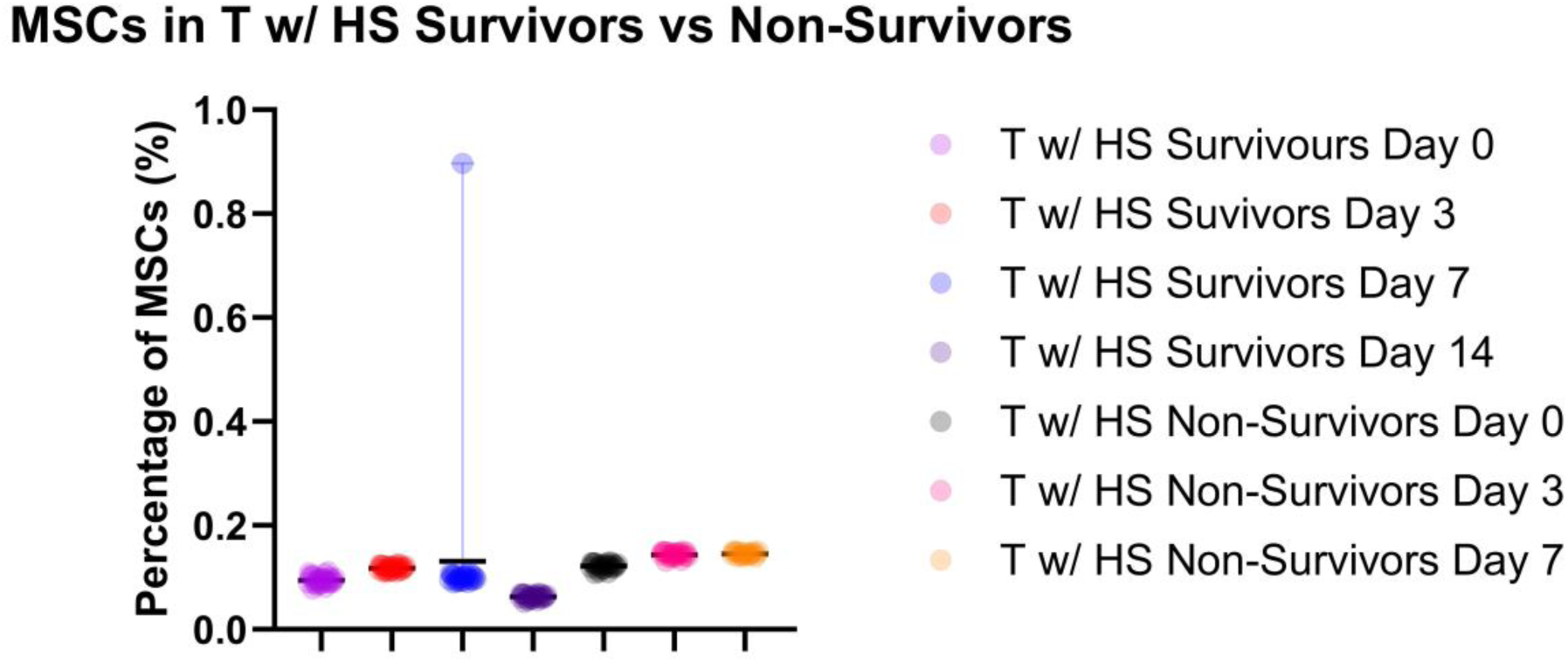
Percentage of Mesenchymal Stromal Cells (MSCs) in Peripheral Human Blood among survivors and non-survivors of hemorrhagic shock, n=50 (Trauma Patients with Hemorrhagic Shock and Traumatic Brain Injury); ****p<0.0001.

**Figure 07:**
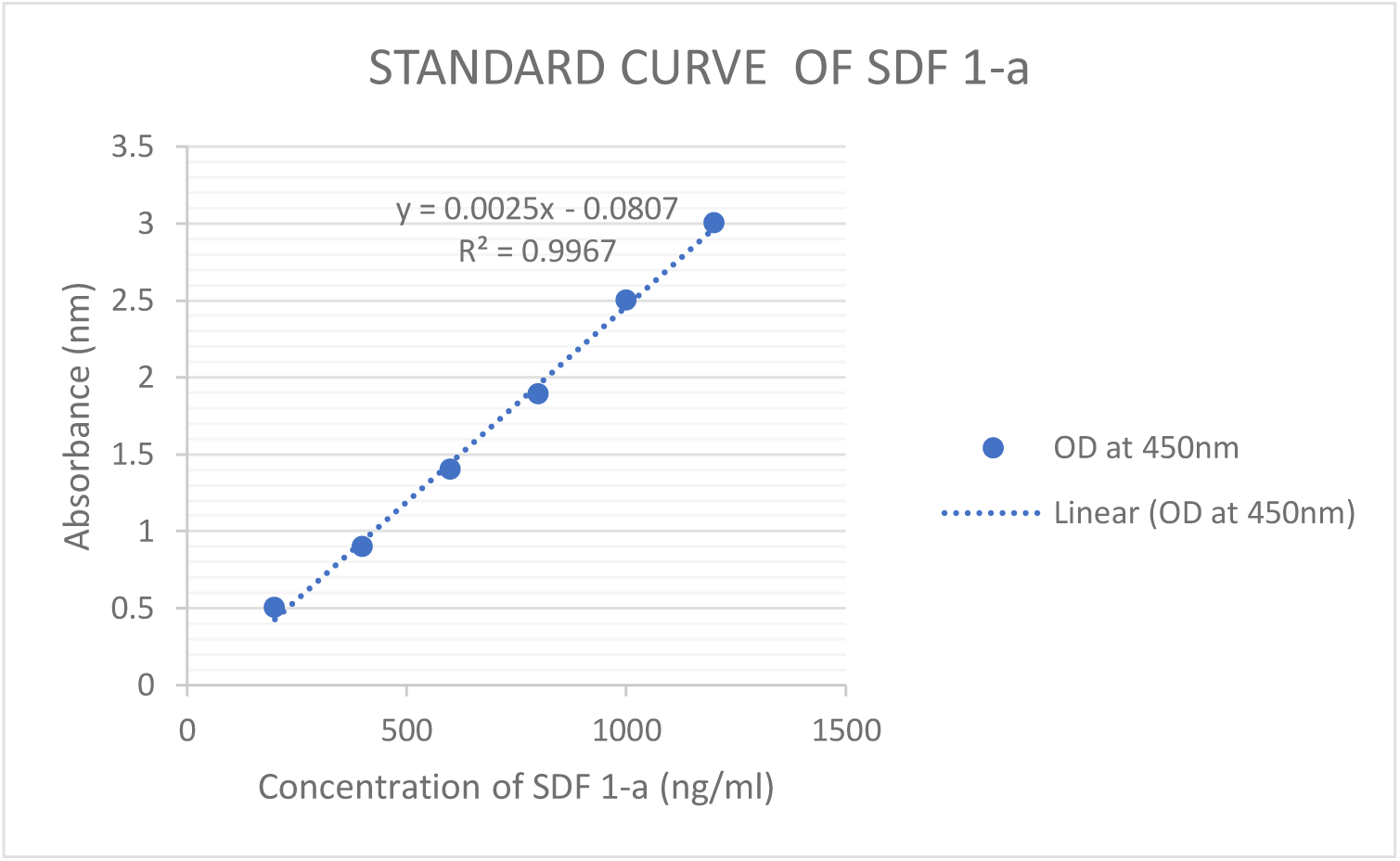
Standard Curve obtained for estimating SDF-1 in the serum sample of comparator group and index group, n=25 (Trauma Patients without Hemorrhagic Shock), n=50 (Trauma Patients with Hemorrhagic Shock and Traumatic Brain Injury).

Thus, persistently high SPC percentages were associated with non-survival, whereas a transient early rise followed by decline was associated with survival.

Cytokine profiles:

All measured cytokines (SDF-1, VEGF-A, EGF, GRO-α, GRO-β, GM-CSF, G-CSF) were significantly elevated in trauma patients without HS at admission compared with minor-injury controls (Figure 08, Figure 12, Figure 16, Figure 20, Figure 24, Figure 28, Figure 32).

**Figure 08:**
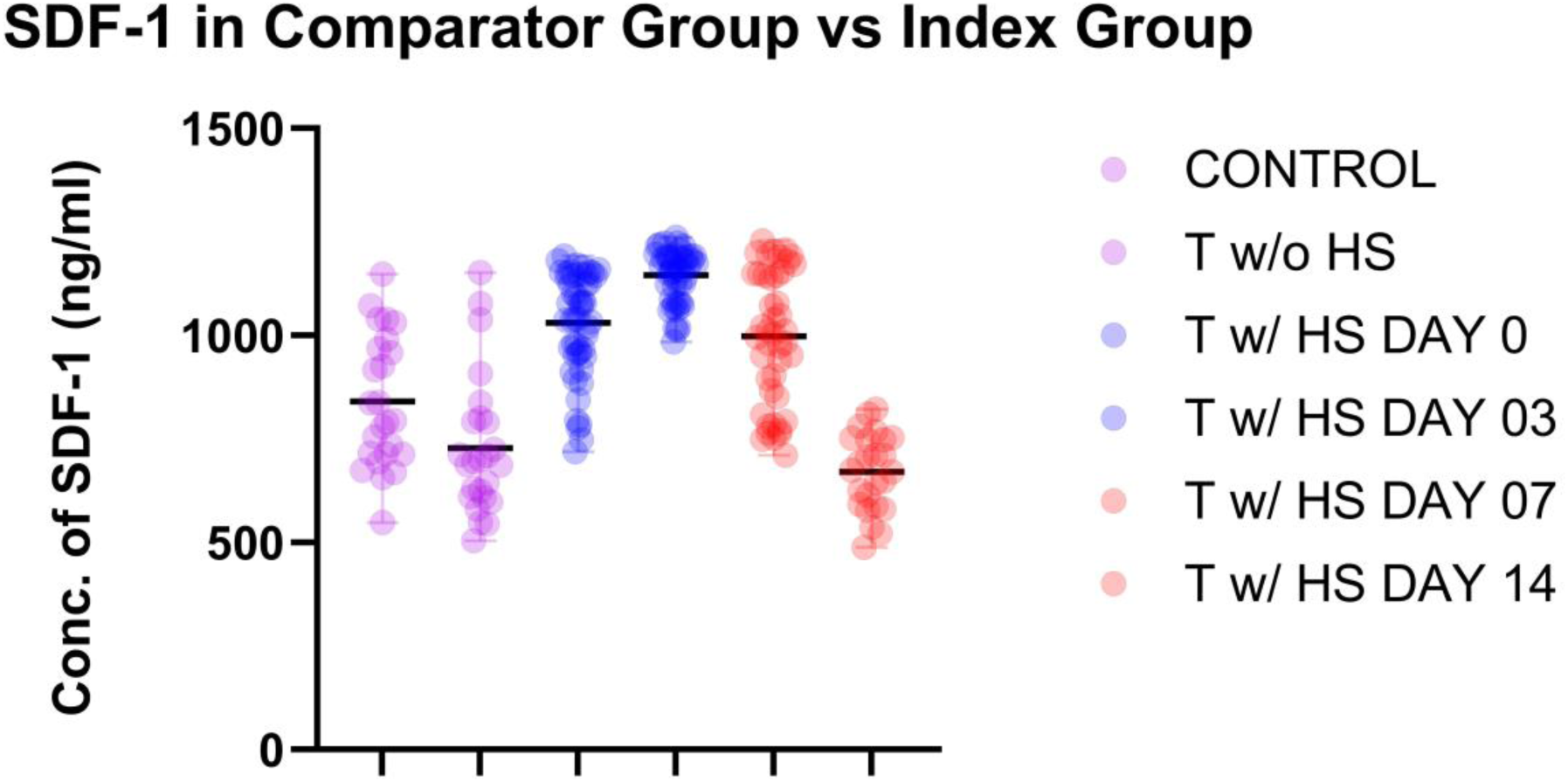
Concentration of SDF-1 in the serum sample of comparator group and index group, n=25 (Trauma Patients without Hemorrhagic Shock), n=50 (Trauma Patients with Hemorrhagic Shock and Traumatic Brain Injury); ****p<0.0001.

In the HS cohort the cytokine time course mirrored SPC kinetics (Figure 09, Figure 13, Figure 17, Figure 21, Figure 25, Figure 29, Figure 33):

- Day 0–3: sharp up-regulation of all cytokines.
- Day 7–14: decline toward lower levels.

**Figure 09:**
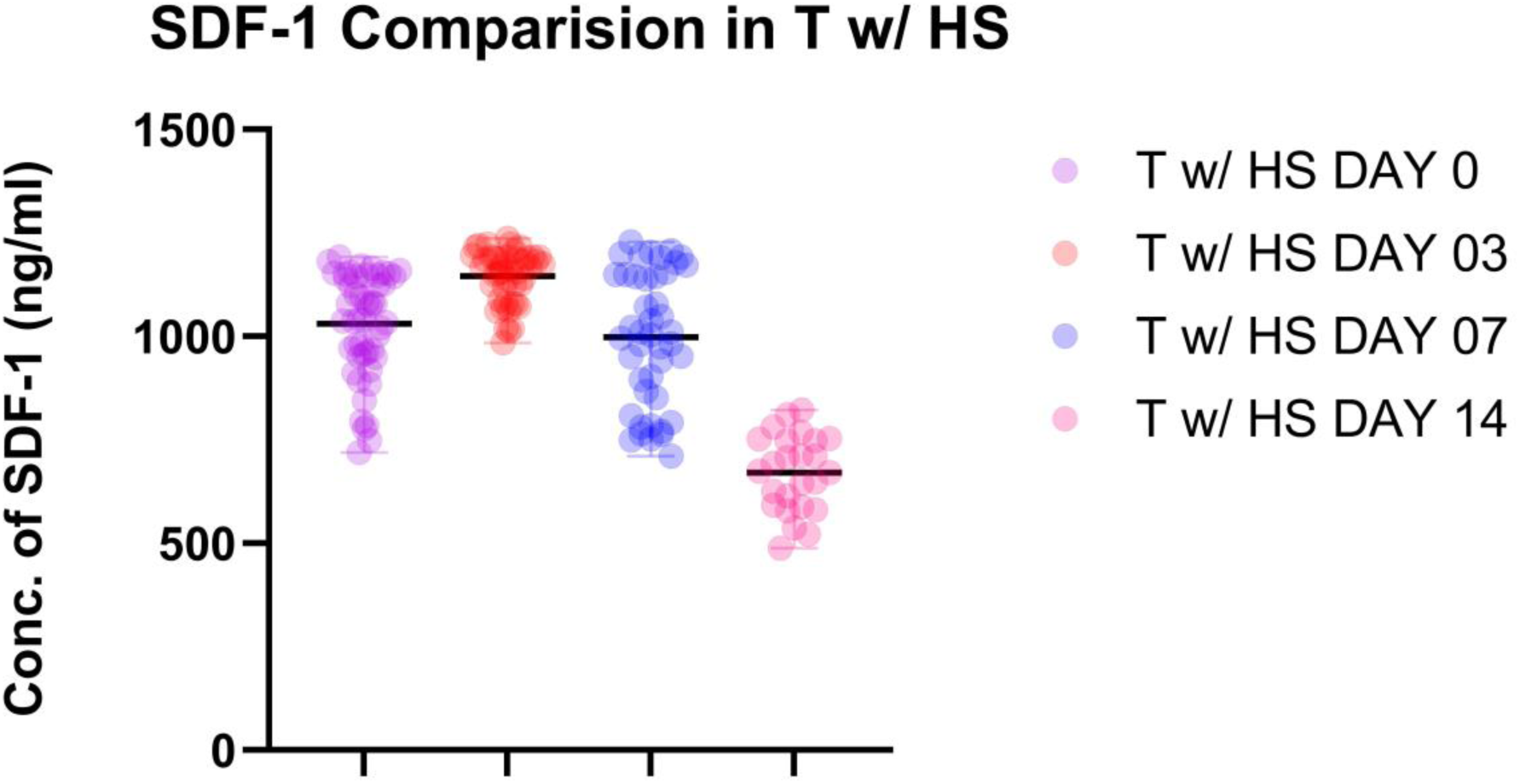
Concentration of SDF-1 in the serum sample of Index Group, n=50 (Trauma Patients with Hemorrhagic Shock and Traumatic Brain Injury); ****p<0.0001.

**Figure 10:**
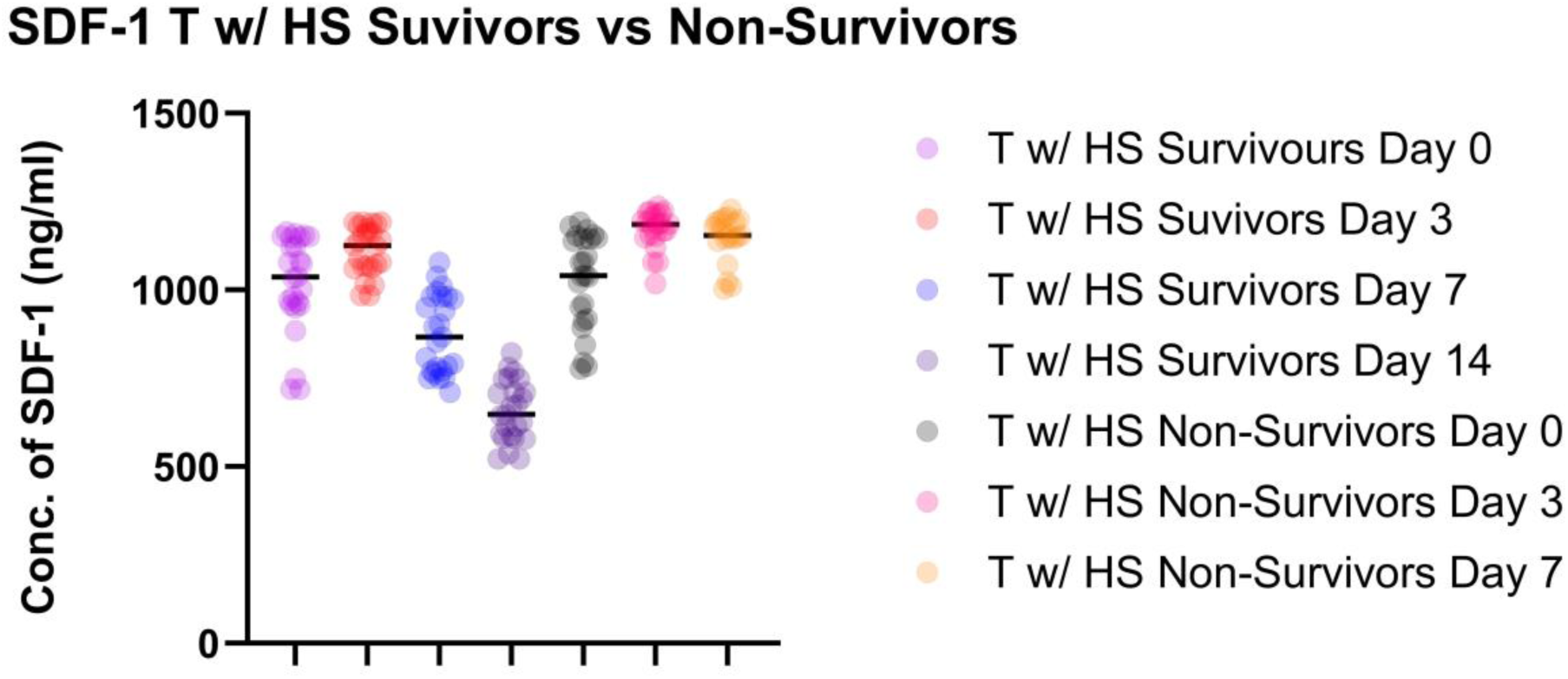
Concentration of SDF-1 in serum samples among survivors and non-survivors of hemorrhagic shock, n=50 (Trauma Patients with Hemorrhagic Shock and Traumatic Brain Injury); ****p<0.0001.

**Figure 11:**
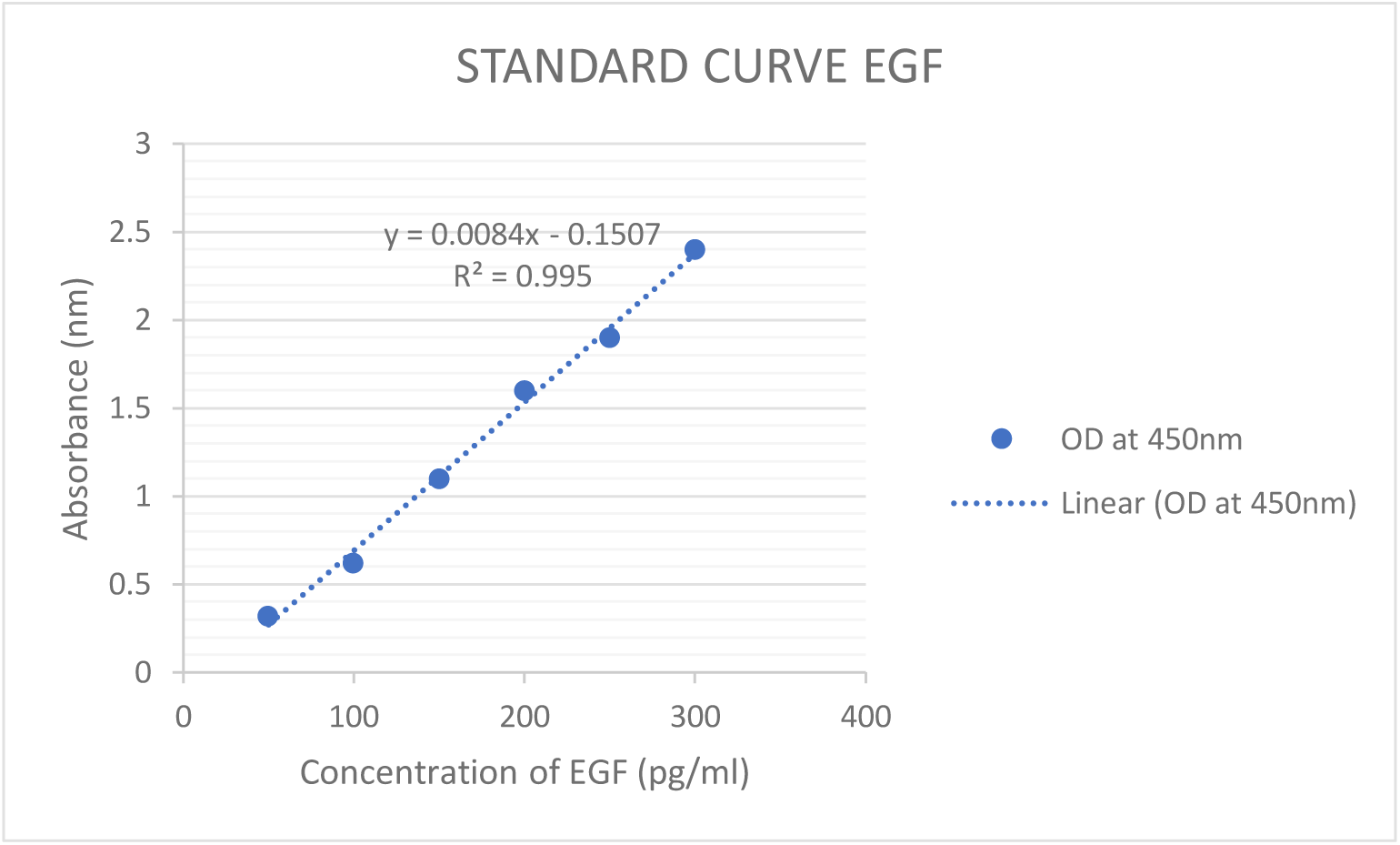
Standard Curve obtained for estimating EGF in the serum sample of comparator group and index group, n=25 (Trauma Patients without Hemorrhagic Shock), n=50 (Trauma Patients with Hemorrhagic Shock and Traumatic Brain Injury).

**Figure 12:**
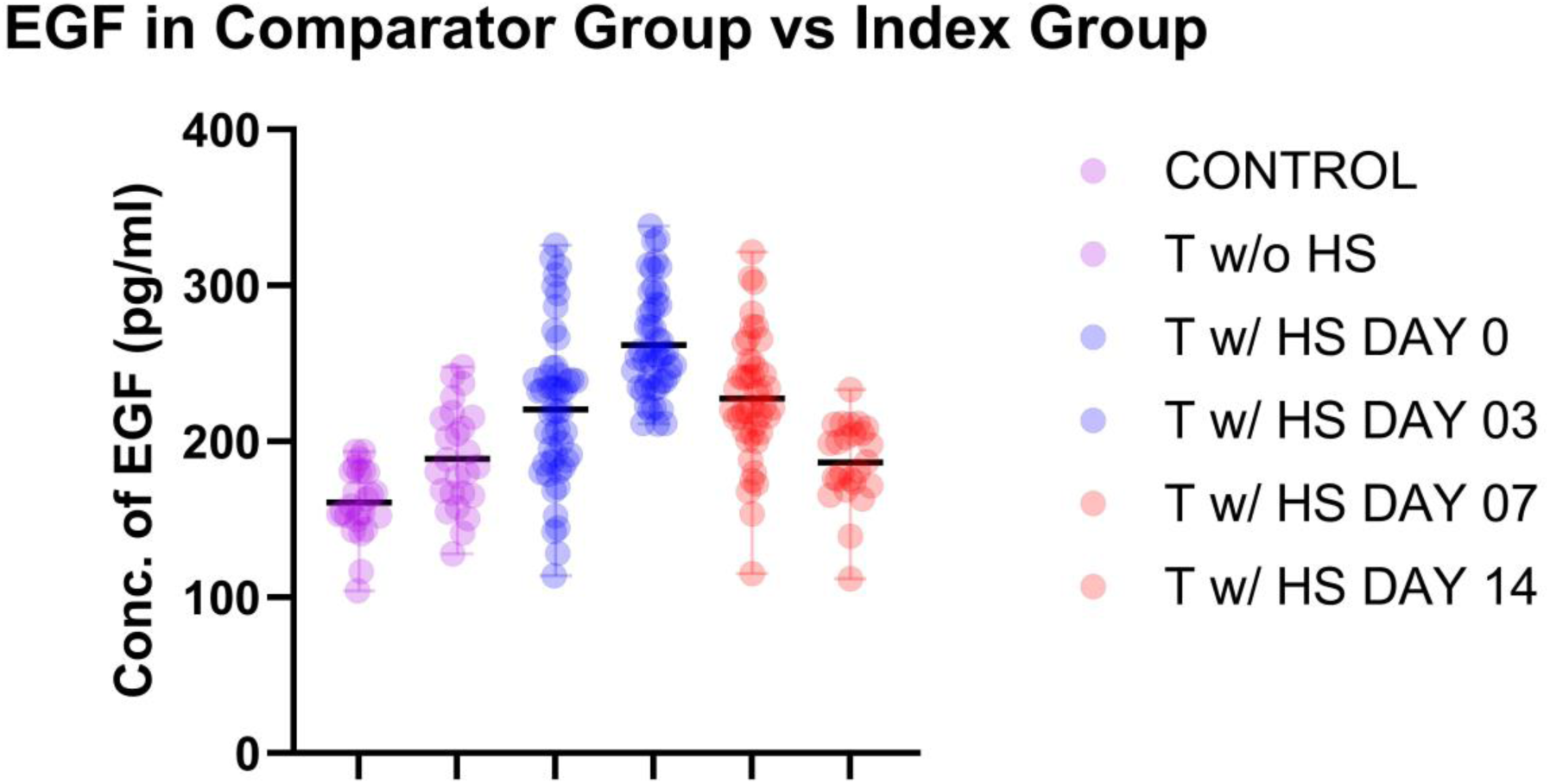
Concentration of EGF in the serum sample of comparator group and index group, n=25 (Trauma Patients without Hemorrhagic Shock), n=50 (Trauma Patients with Hemorrhagic Shock and Traumatic Brain Injury); ****p<0.0001.

**Figure 13:**
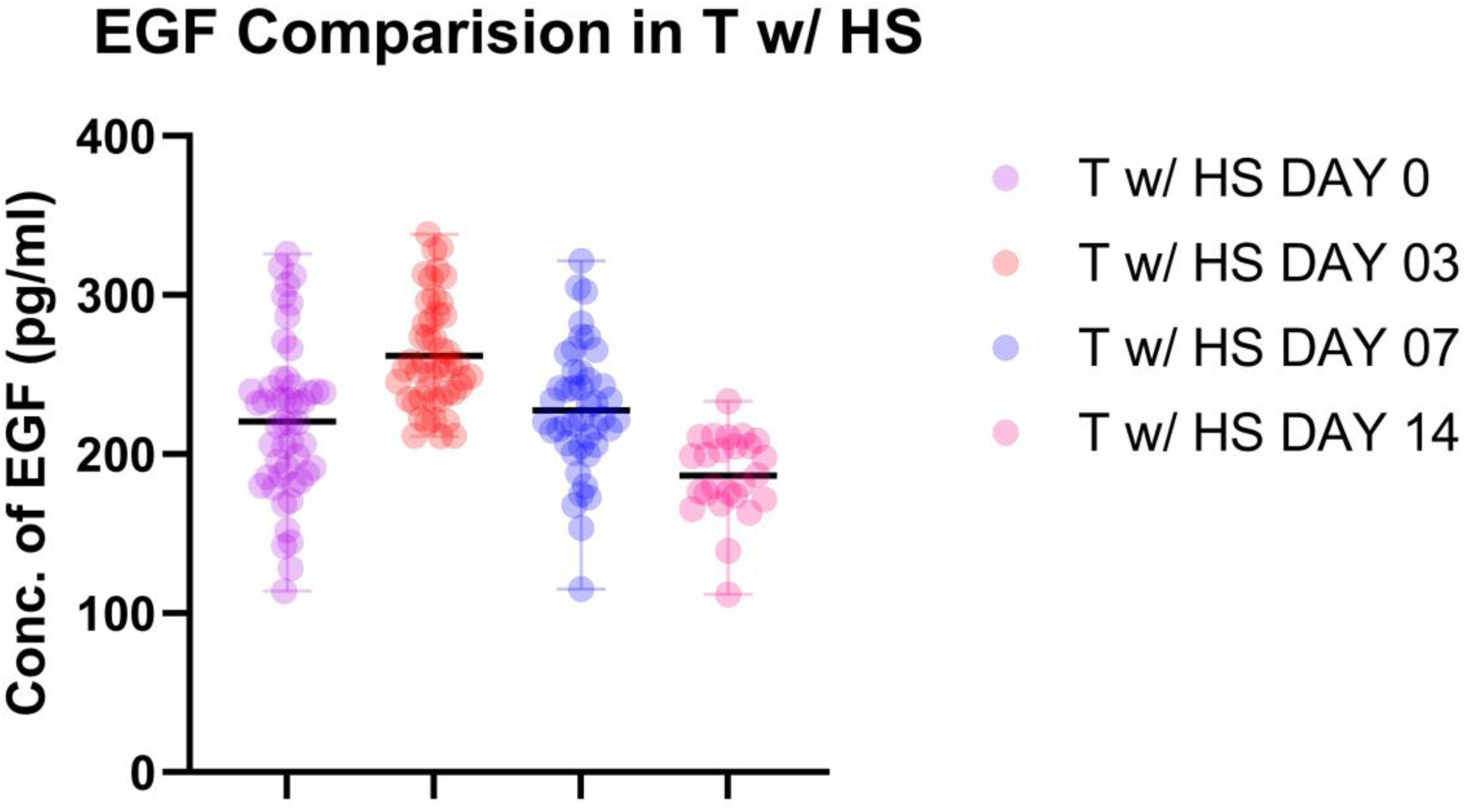
Concentration of EGF in the serum sample of Index Group, n=50 (Trauma Patients with Hemorrhagic Shock and Traumatic Brain Injury); ****p<0.0001.

Non-survivors exhibited higher cytokine concentrations than survivors at comparable time points; elevated cytokines persisted in non-survivors up to death (Figure 10, Figure 14, Figure 18, Figure 22, Figure 26, Figure 30, Figure 34).

**Figure 14:**
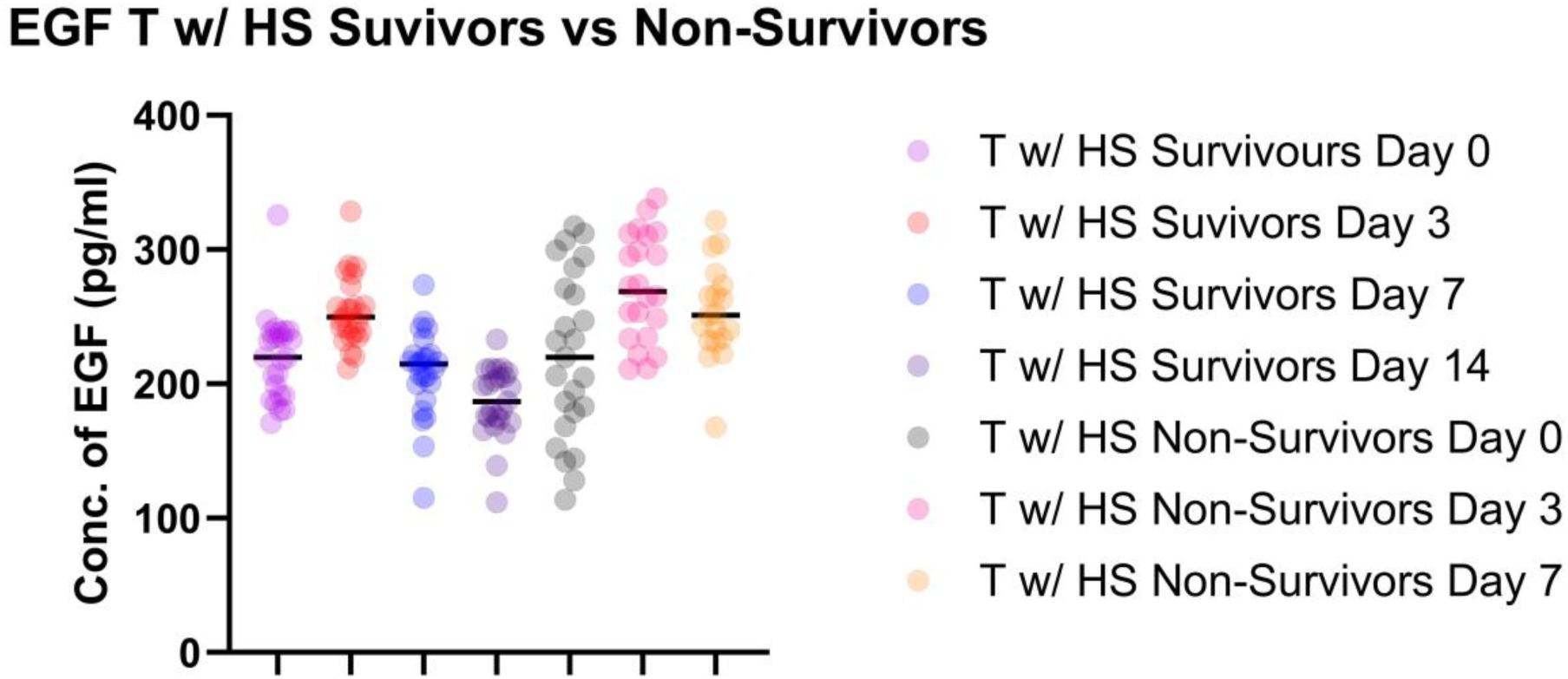
Concentration of EGF in serum samples among survivors and non-survivors of hemorrhagic shock, n=50 (Trauma Patients with Hemorrhagic Shock and Traumatic Brain Injury); ****p<0.0001

**Figure 15:**
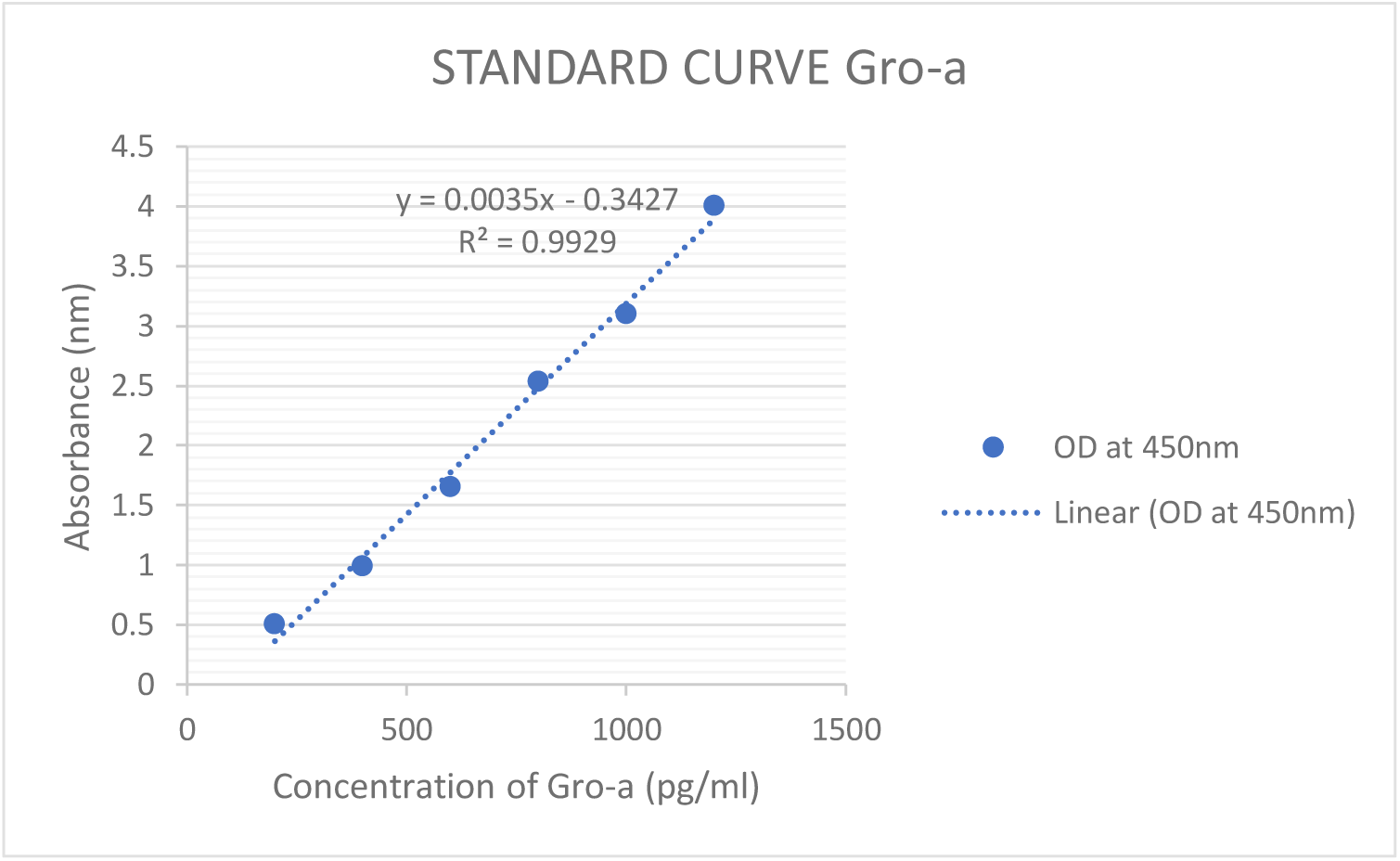
Standard Curve obtained for estimating GRO-A in the serum sample of comparator group and index group, n=25 (Trauma Patients without Hemorrhagic Shock), n=50 (Trauma Patients with Hemorrhagic Shock and Traumatic Brain Injury).

**Figure 16:**
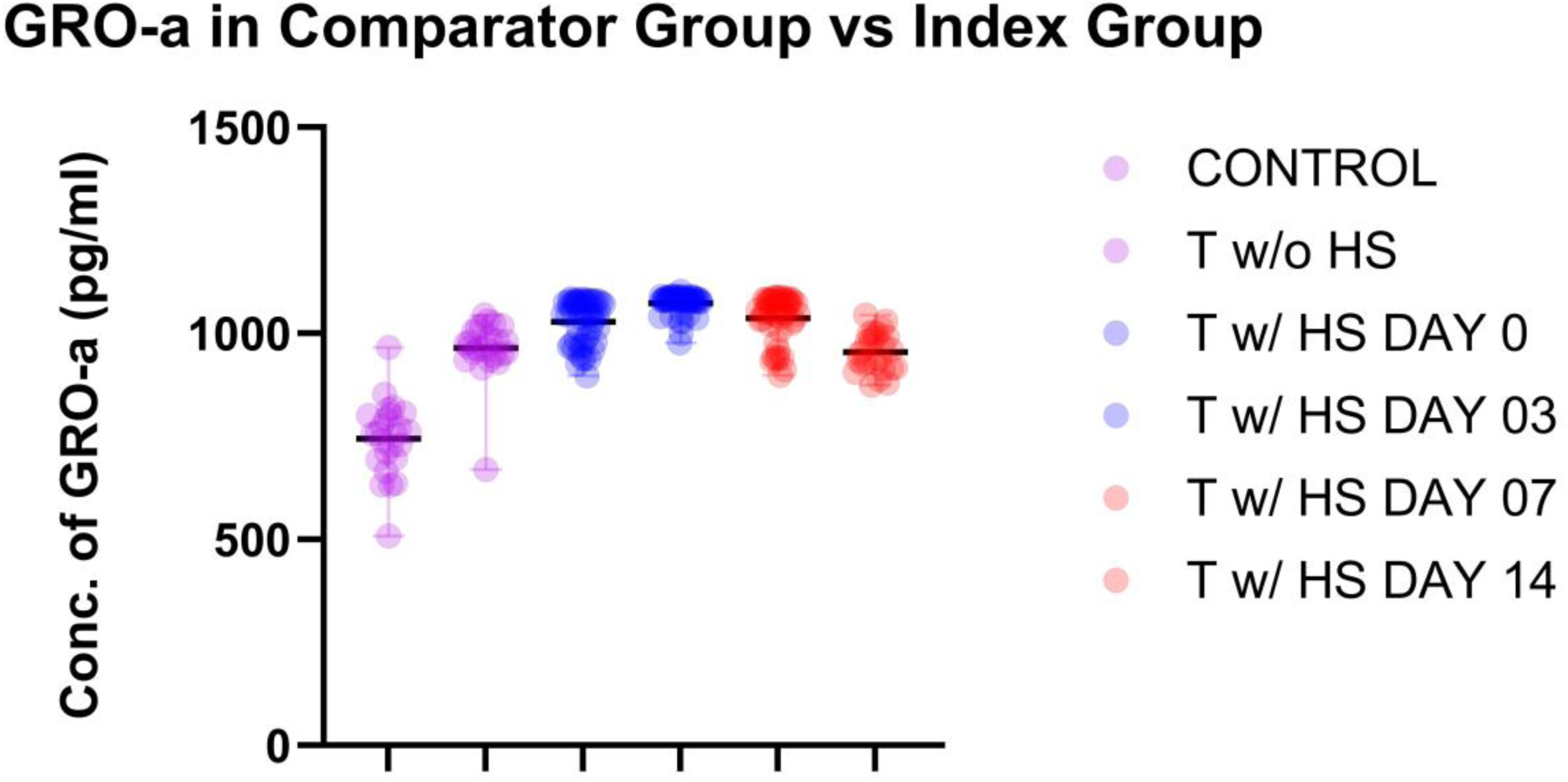
Concentration of GRO-A in the serum sample of comparator group and index group, n=25 (Trauma Patients without Hemorrhagic Shock), n=50 (Trauma Patients with Hemorrhagic Shock and Traumatic Brain Injury); ****p<0.0001.

**Figure 17:**
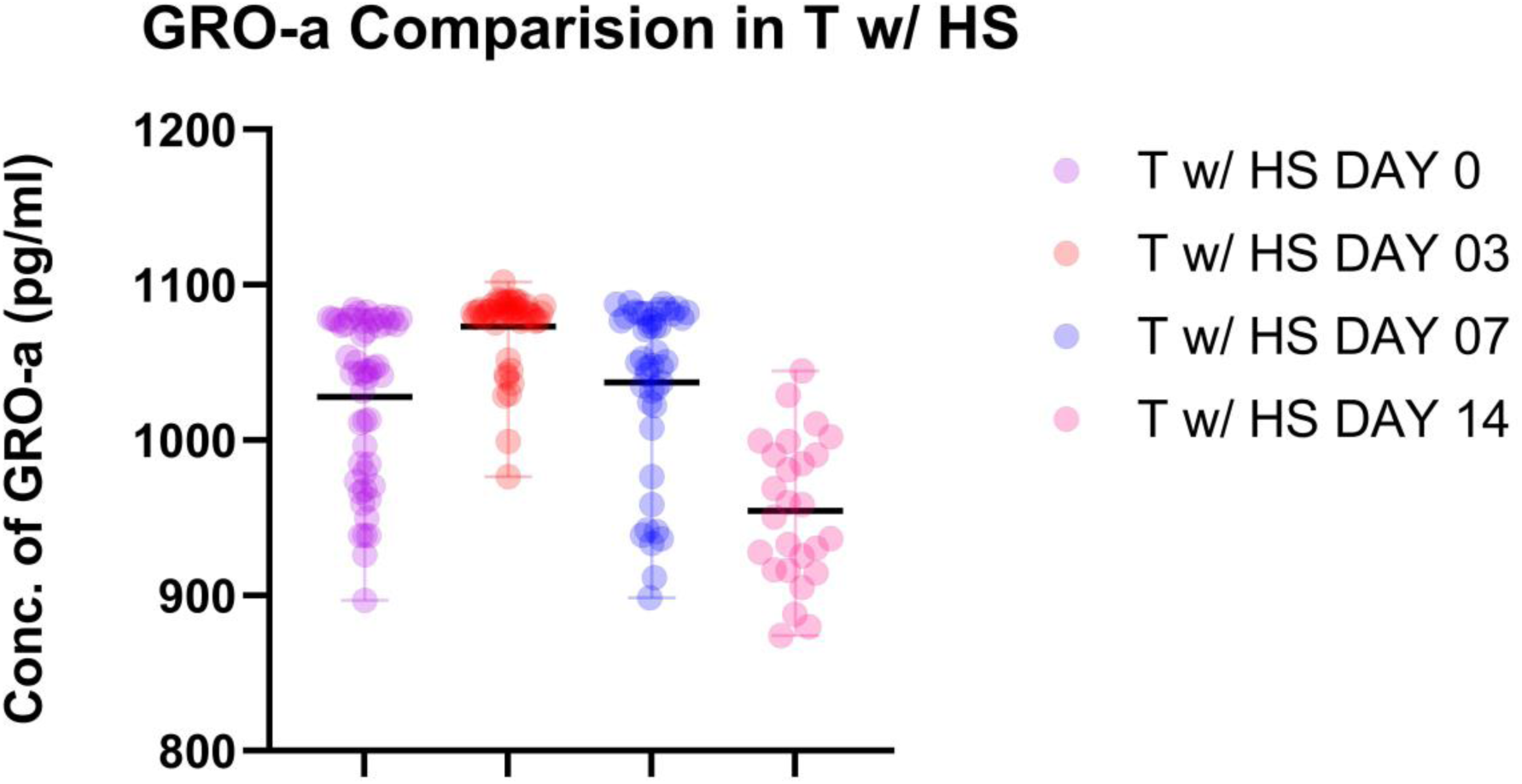
Concentration of GRO-A in the serum sample of Index Group, n=50 (Trauma Patients with Hemorrhagic Shock and Traumatic Brain Injury); ****p<0.0001.

**Figure 18:**
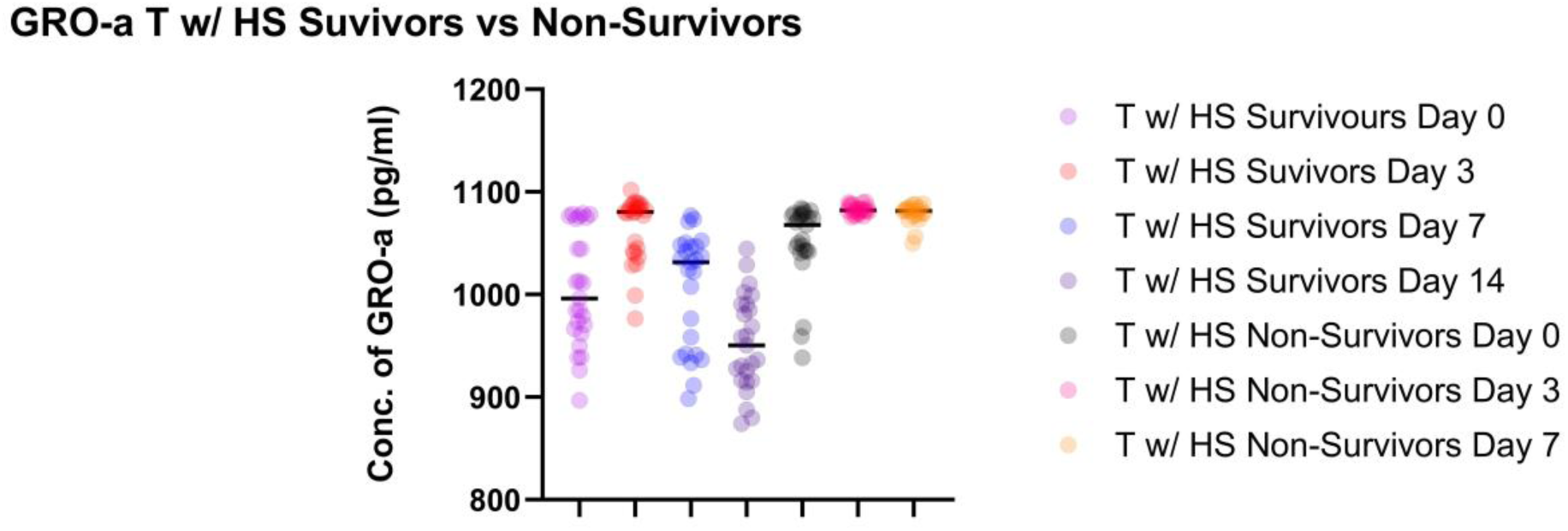
Concentration of GRO-A in serum samples among survivors and non-survivors of hemorrhagic shock, n=50 (Trauma Patients with Hemorrhagic Shock and Traumatic Brain Injury); ****p<0.0001

**Figure 19:**
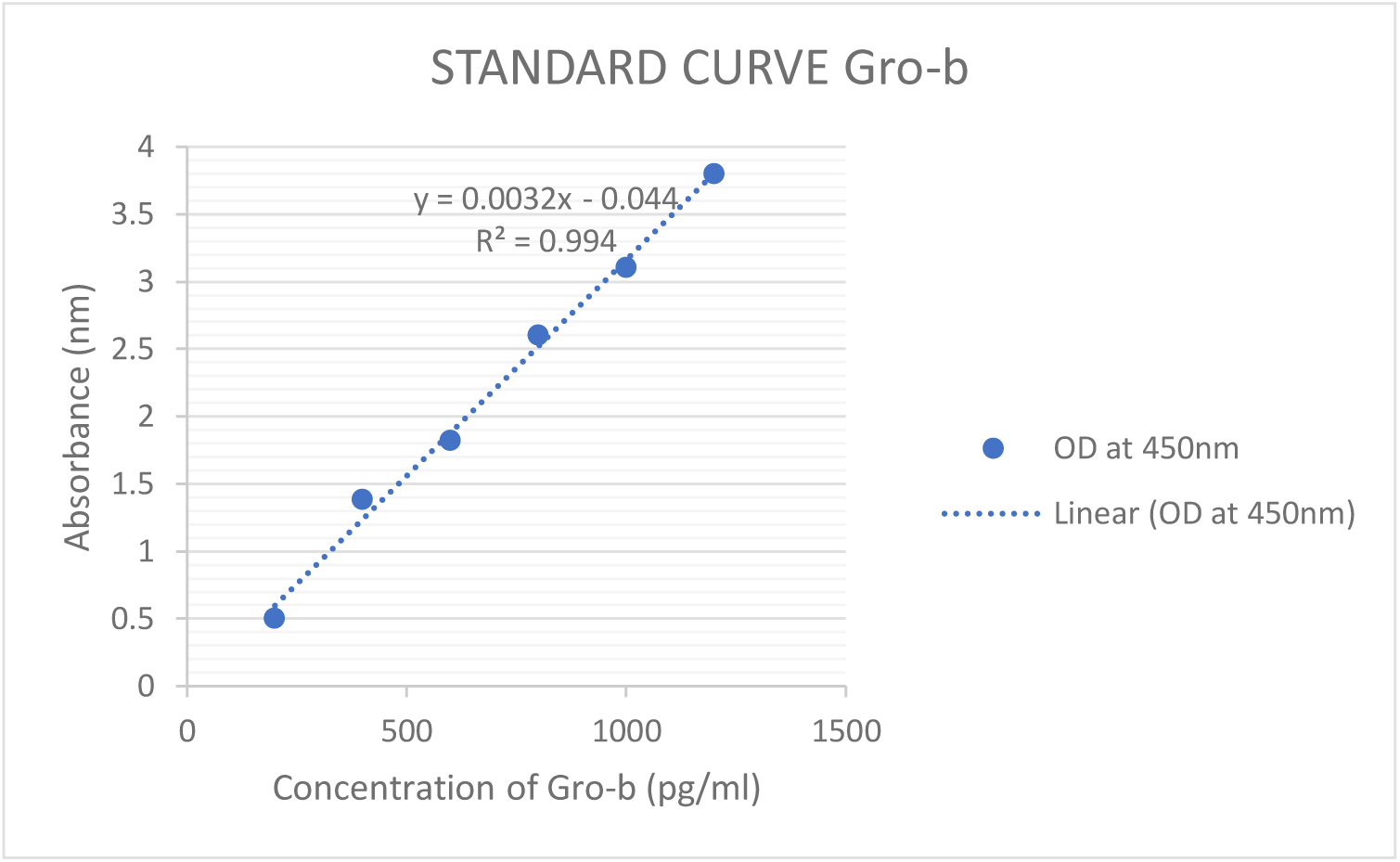
Standard Curve obtained for estimating GRO-B in the serum sample of comparator group and index group, n=25 (Trauma Patients without Hemorrhagic Shock), n=50 (Trauma Patients with Hemorrhagic Shock and Traumatic Brain Injury).

**Figure 20:**
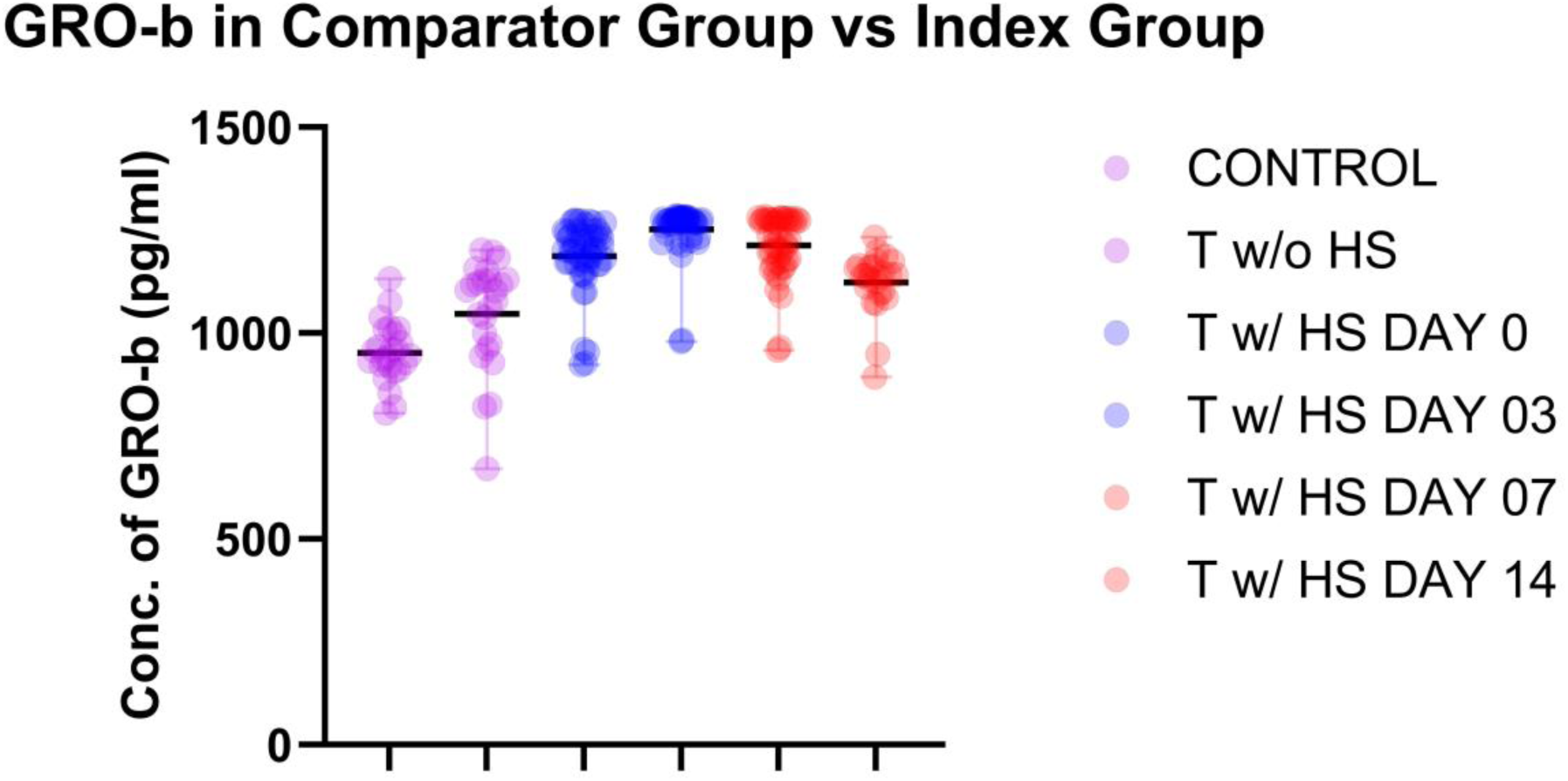
Concentration of GRO-B in the serum sample of comparator group and index group, n=25 (Trauma Patients without Hemorrhagic Shock), n=50 (Trauma Patients with Hemorrhagic Shock and Traumatic Brain Injury); **** p<0.0001.

**Figure 21:**
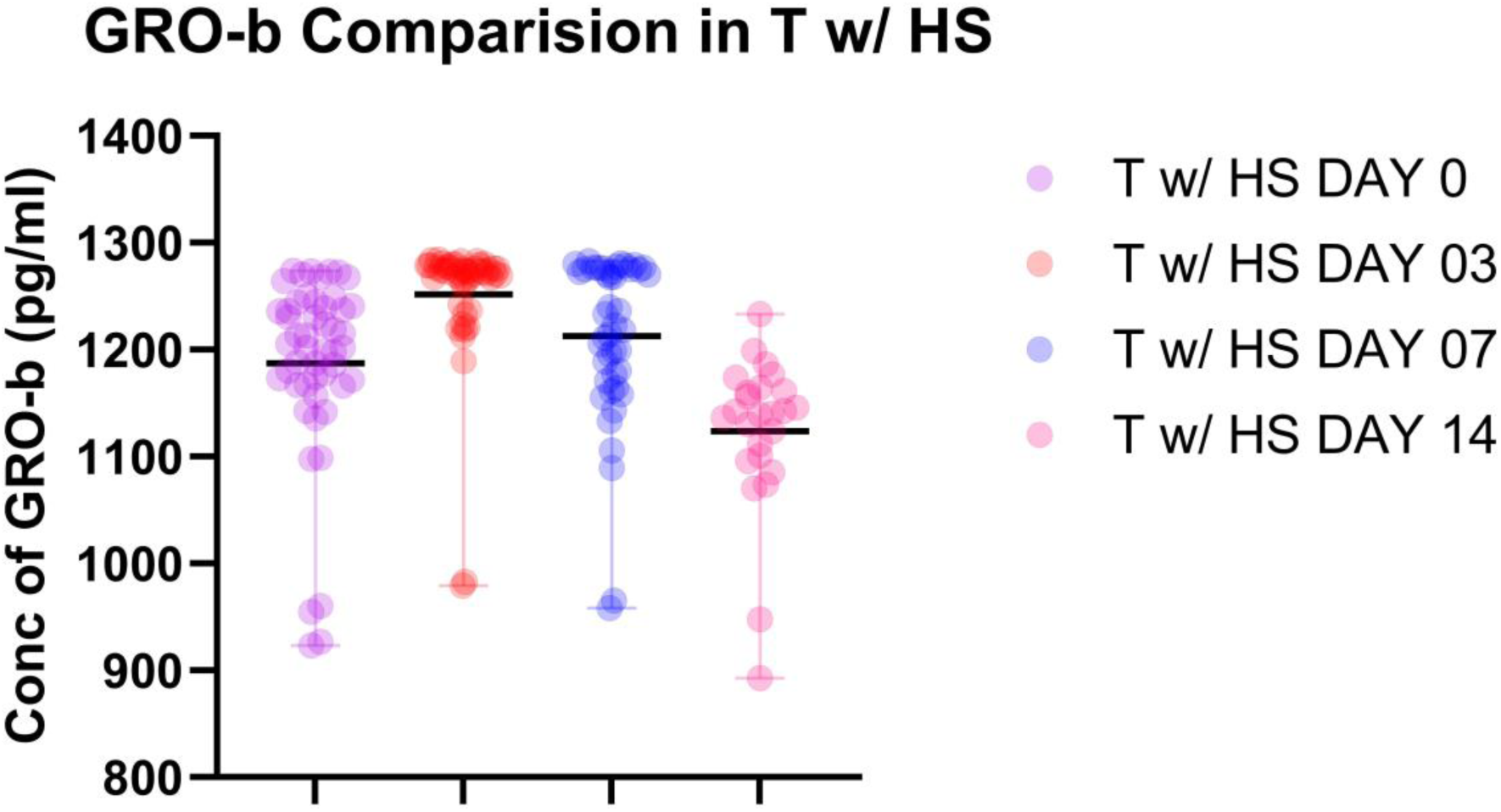
Concentration of GRO-B in the serum sample of Index Group, n=50 (Trauma Patients with Hemorrhagic Shock and Traumatic Brain Injury); ****p<0.0001.

**Figure 22:**
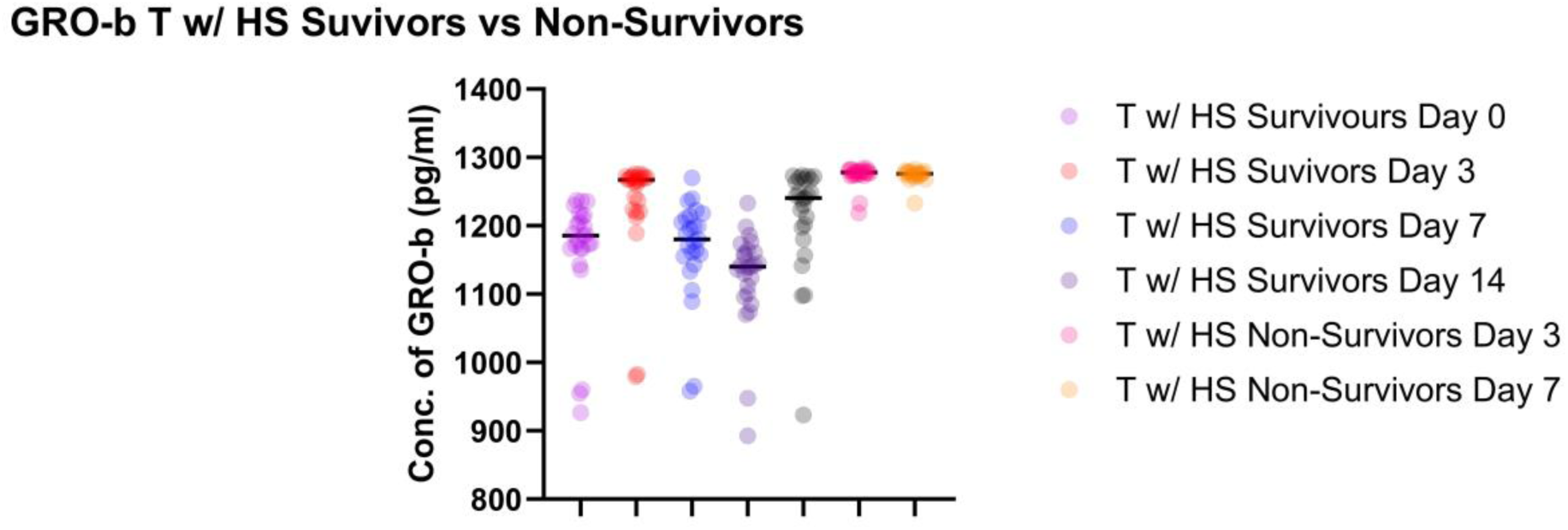
Concentration of GRO-B in serum samples among survivors and non-survivors of hemorrhagic shock, n=50 (Trauma Patients with Hemorrhagic Shock and Traumatic Brain Injury); ****p<0.0001.

**Figure 23:**
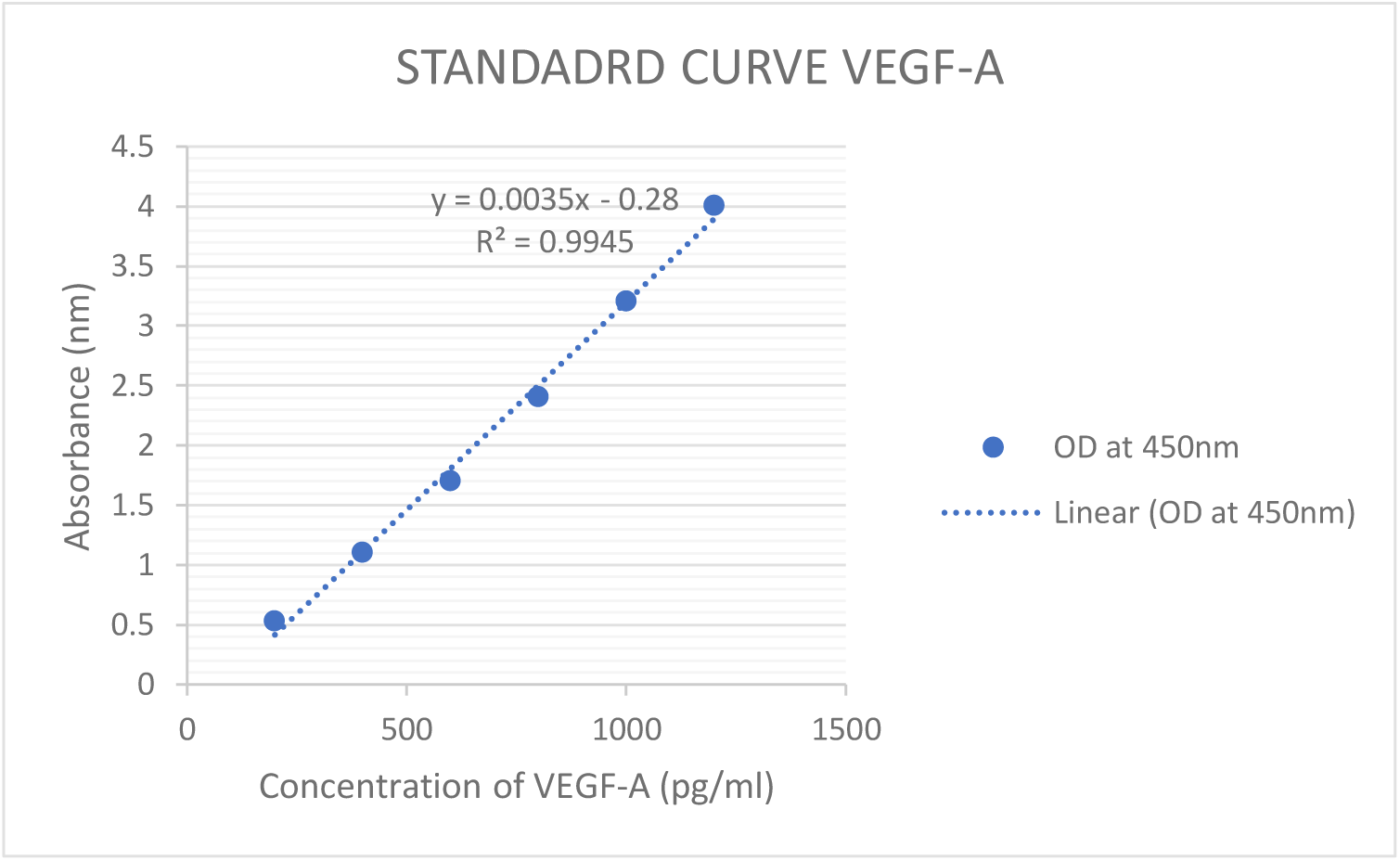
Standard Curve obtained for estimating VEGF-A in the serum sample of comparator group and index group, n=25 (Trauma Patients without Hemorrhagic Shock), n=50 (Trauma Patients with Hemorrhagic Shock and Traumatic Brain Injury).

**Figure 24:**
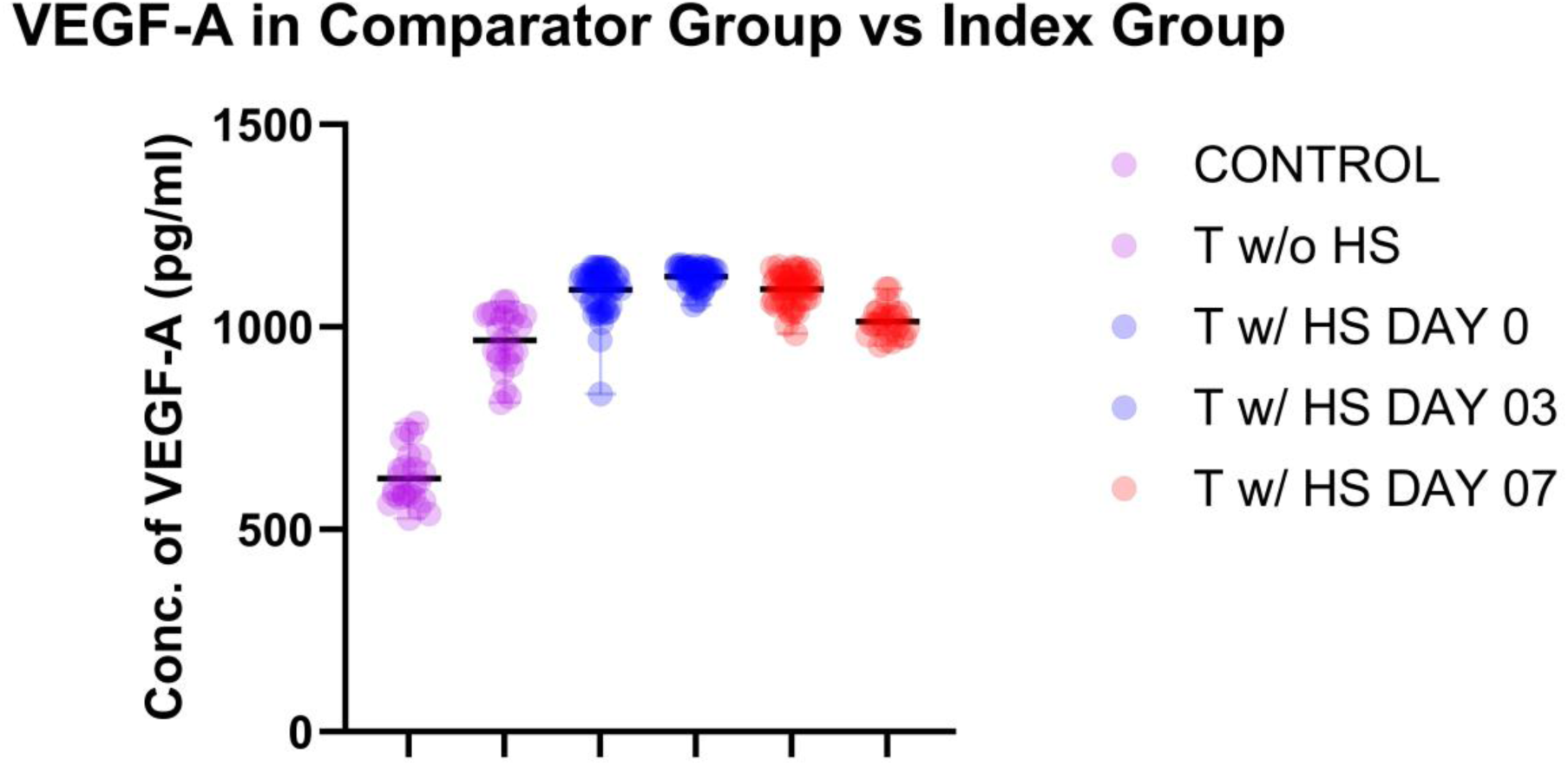
Concentration of VEGF-A in the serum sample of comparator group and index group, n=25 (Trauma Patients without Hemorrhagic Shock), n=50 (Trauma Patients with Hemorrhagic Shock and Traumatic Brain Injury); ****p<0.0001.

**Figure 25:**
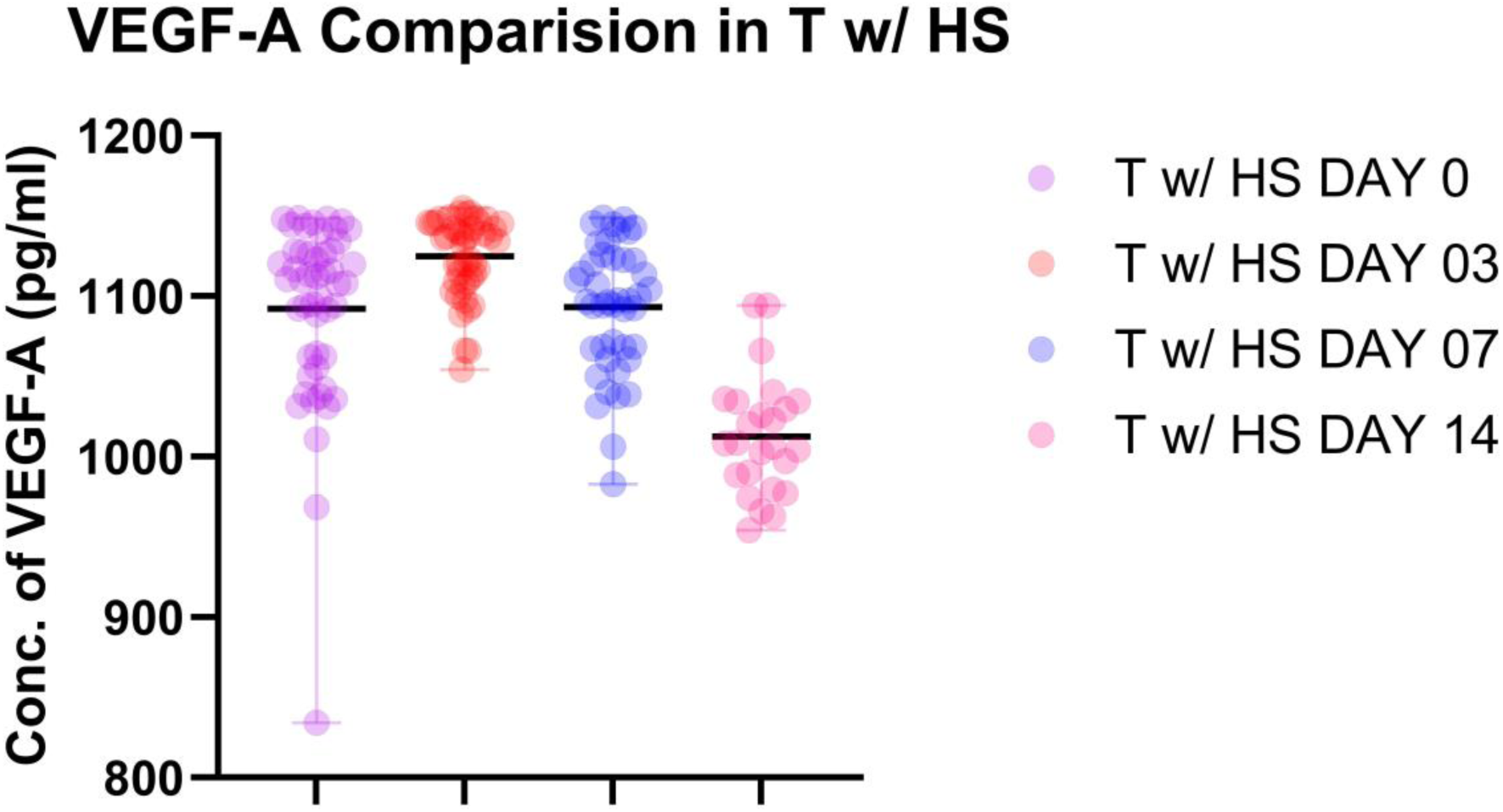
Concentration of VEGF-A in the serum sample of Index Group, n=50 (Trauma Patients with Hemorrhagic Shock and Traumatic Brain Injury); ****p<0.0001.

**Figure 26:**
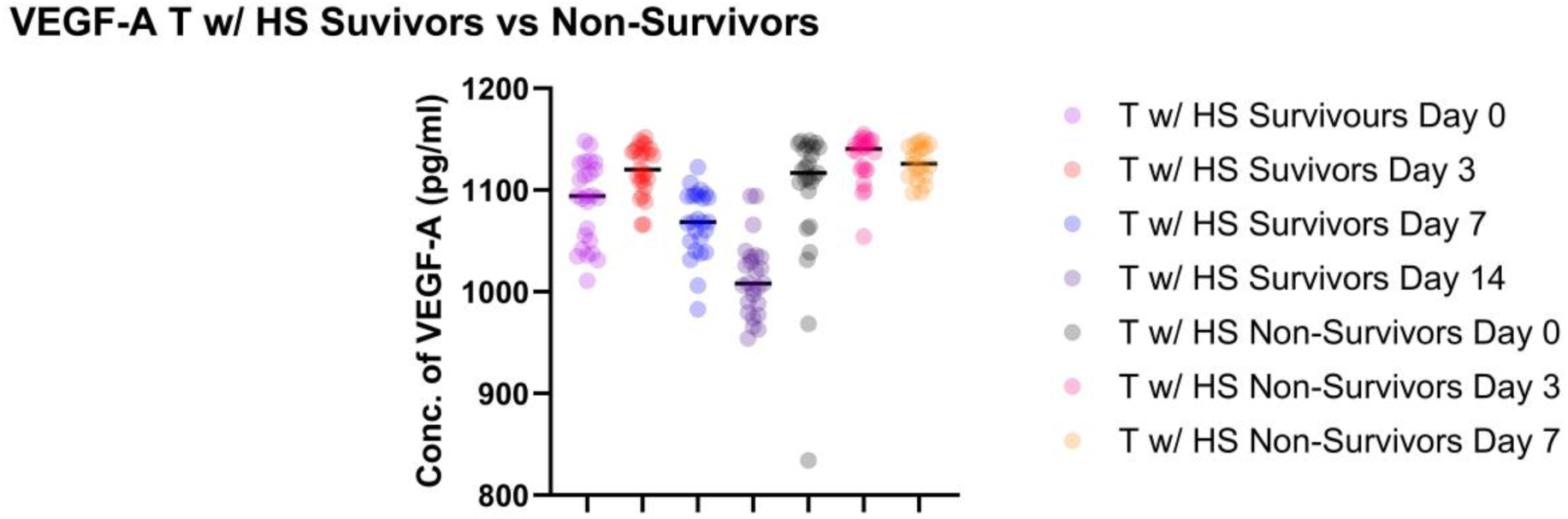
Concentration of VEGF-A in serum samples among survivors and non-survivors of hemorrhagic shock, n=50 (Trauma Patients with Hemorrhagic Shock and Traumatic Brain Injury); ****p<0.0001.

**Figure 27:**
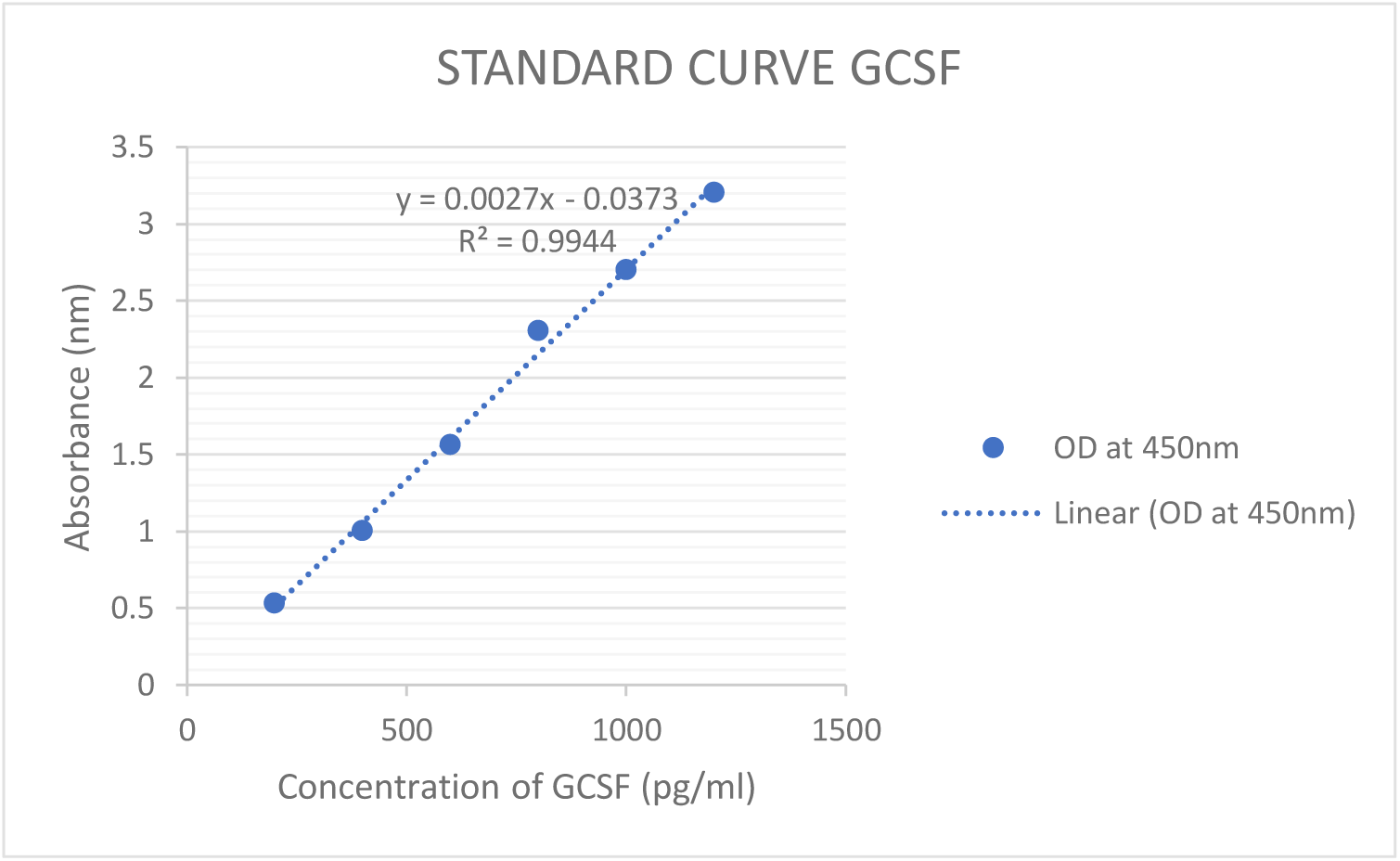
Standard Curve obtained for estimating GCSF in the serum sample of comparator group and index group, n=25 (Trauma Patients without Hemorrhagic Shock), n=50 (Trauma Patients with Hemorrhagic Shock and Traumatic Brain Injury).

**Figure 28:**
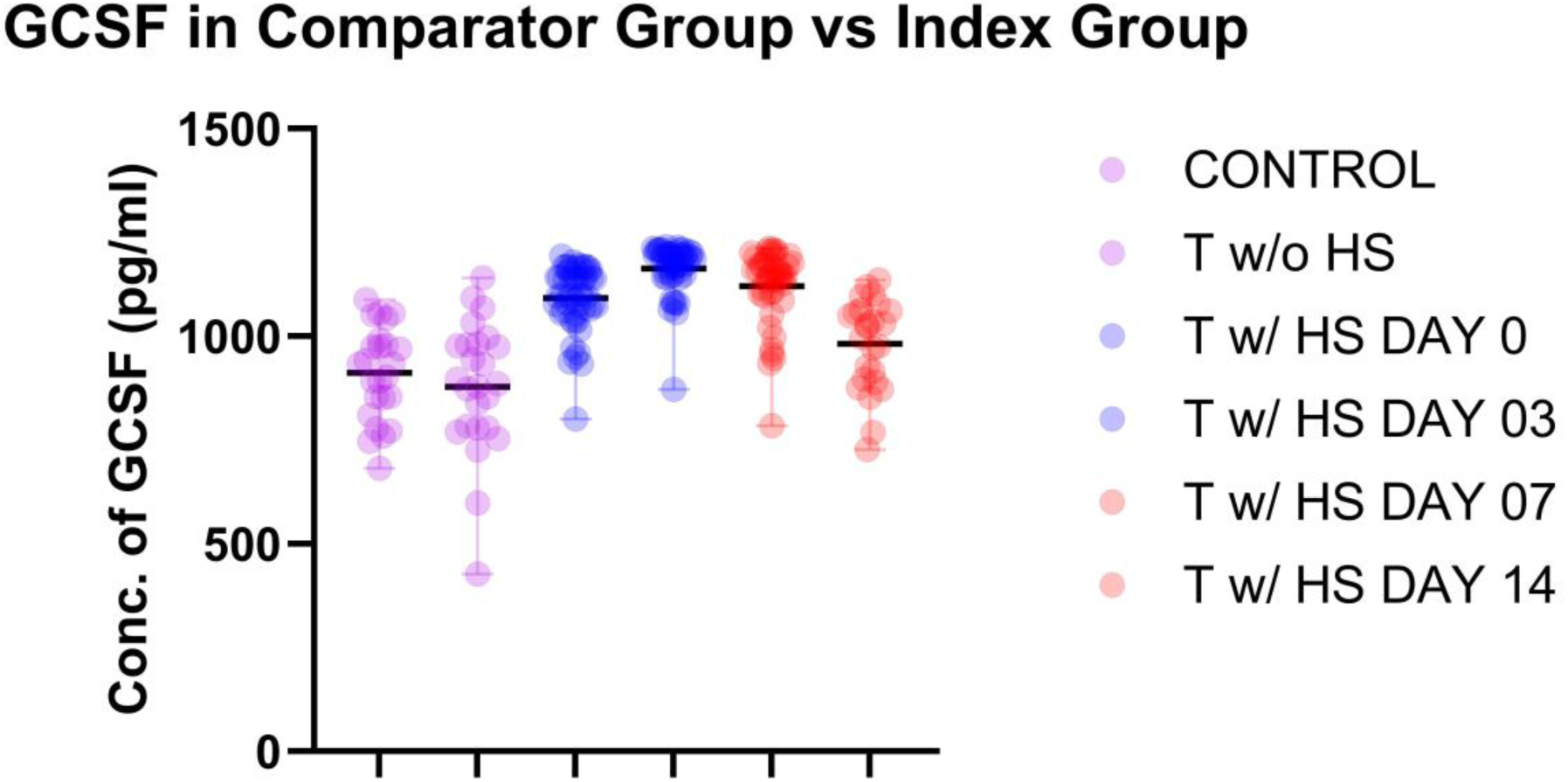
Concentration of GCSF in the serum sample of comparator group and index group, n=25 (Trauma Patients without Hemorrhagic Shock), n=50 (Trauma Patients with Hemorrhagic Shock and Traumatic Brain Injury); ****p<0.0001.

**Figure 29:**
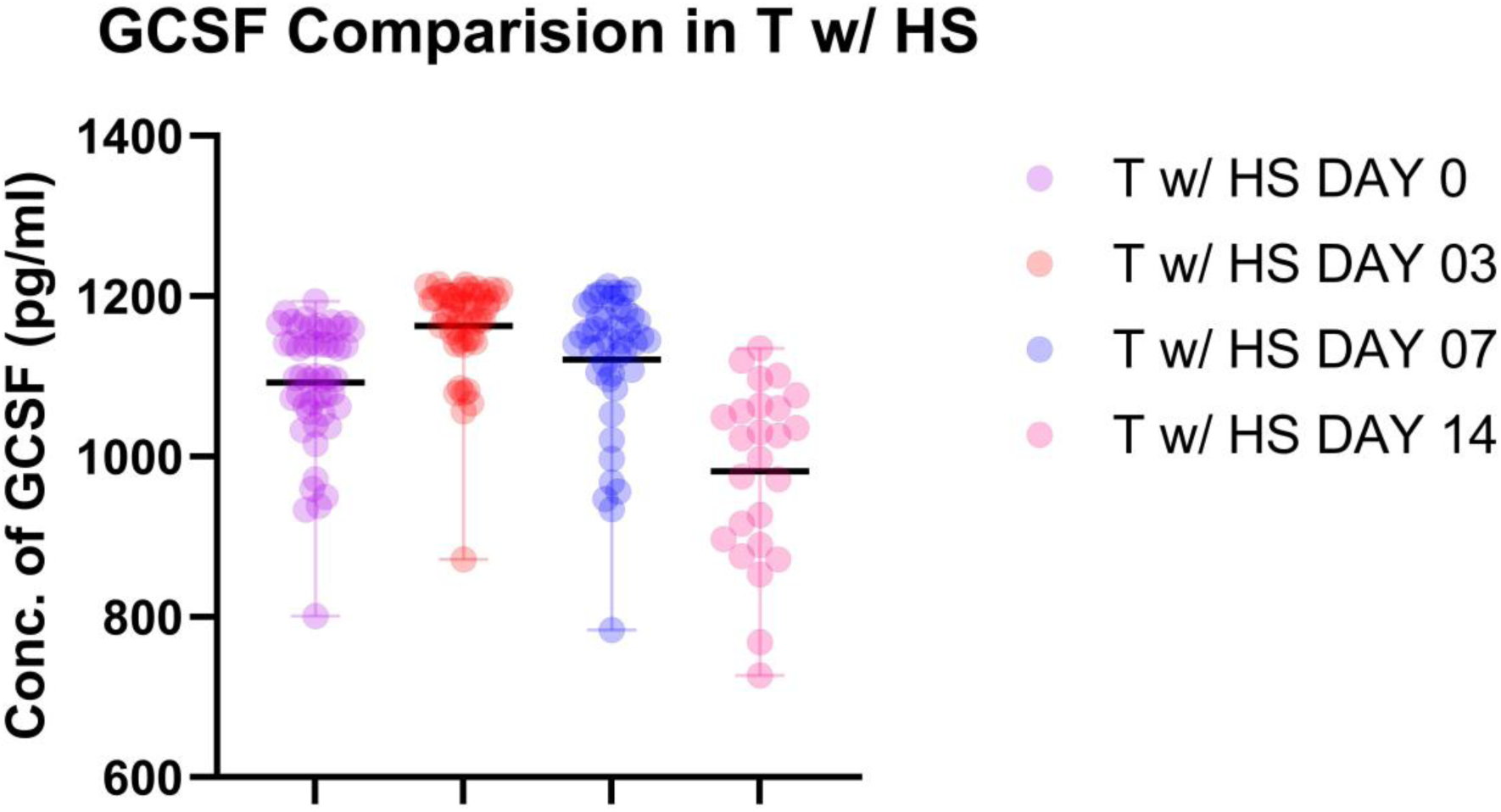
Concentration of GCSF in the serum sample of Index Group, n=50 (Trauma Patients with Hemorrhagic Shock and Traumatic Brain Injury); ****p<0.0001.

**Figure 30:**
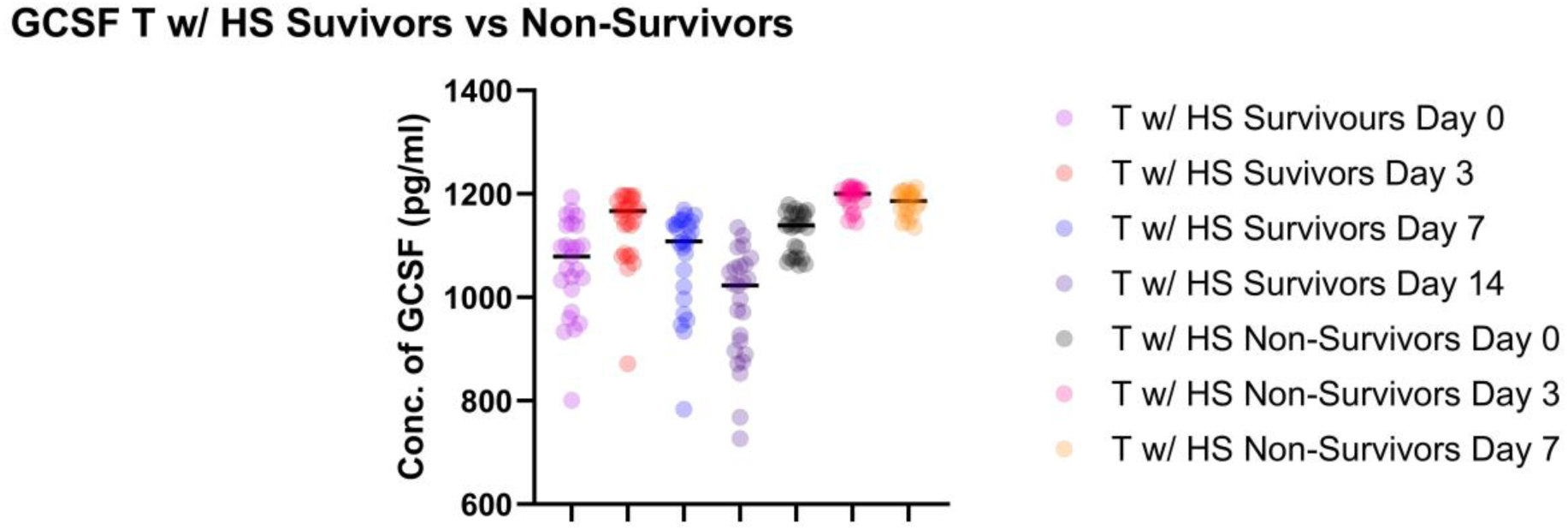
Concentration of GCSF in serum samples among survivors and non-survivors of hemorrhagic shock, n=50 (Trauma Patients with Hemorrhagic Shock and Traumatic Brain Injury); ****p<0.0001.

**Figure 31:**
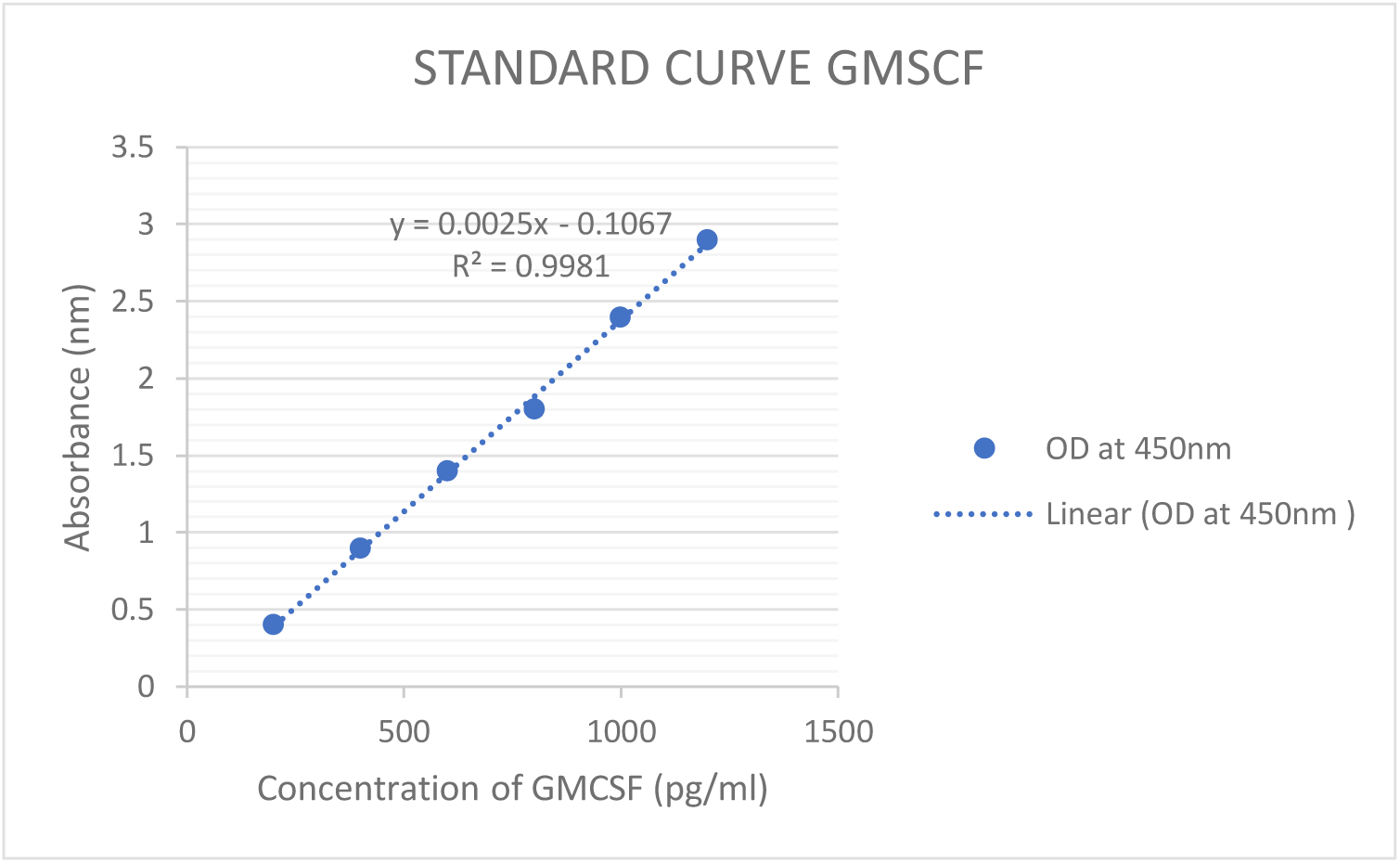
Standard Curve obtained for estimating GMCSF in the serum sample of comparator group and index group, n=25 (Trauma Patients without Hemorrhagic Shock), n=50 (Trauma Patients with Hemorrhagic Shock and Traumatic Brain Injury).

**Figure 32:**
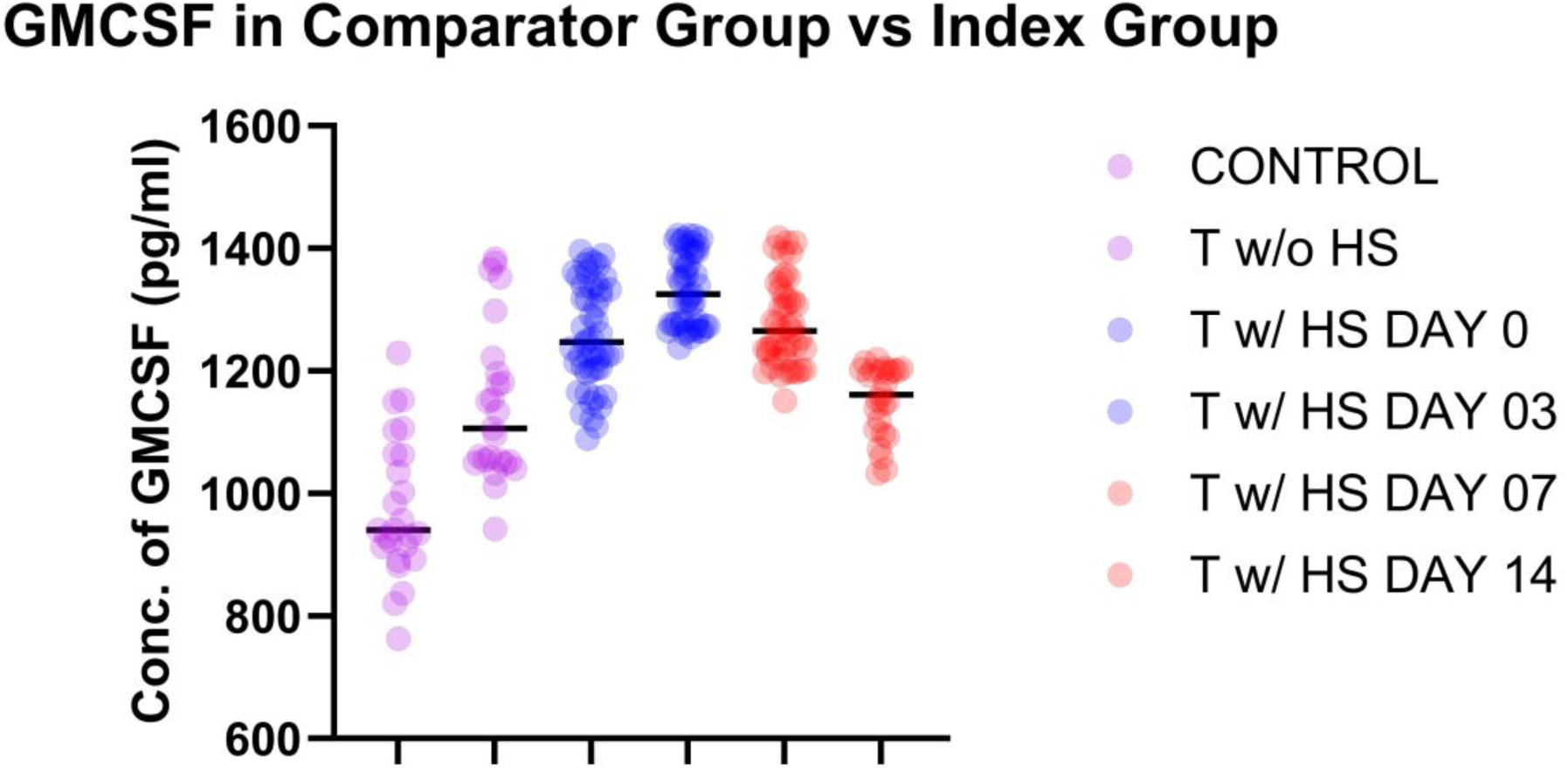
Concentration of GMCSF in the serum sample of comparator group and index group, n=25 (Trauma Patients without Hemorrhagic Shock), n=50 (Trauma Patients with Hemorrhagic Shock and Traumatic Brain Injury); ****p<0.0001.

**Figure 33:**
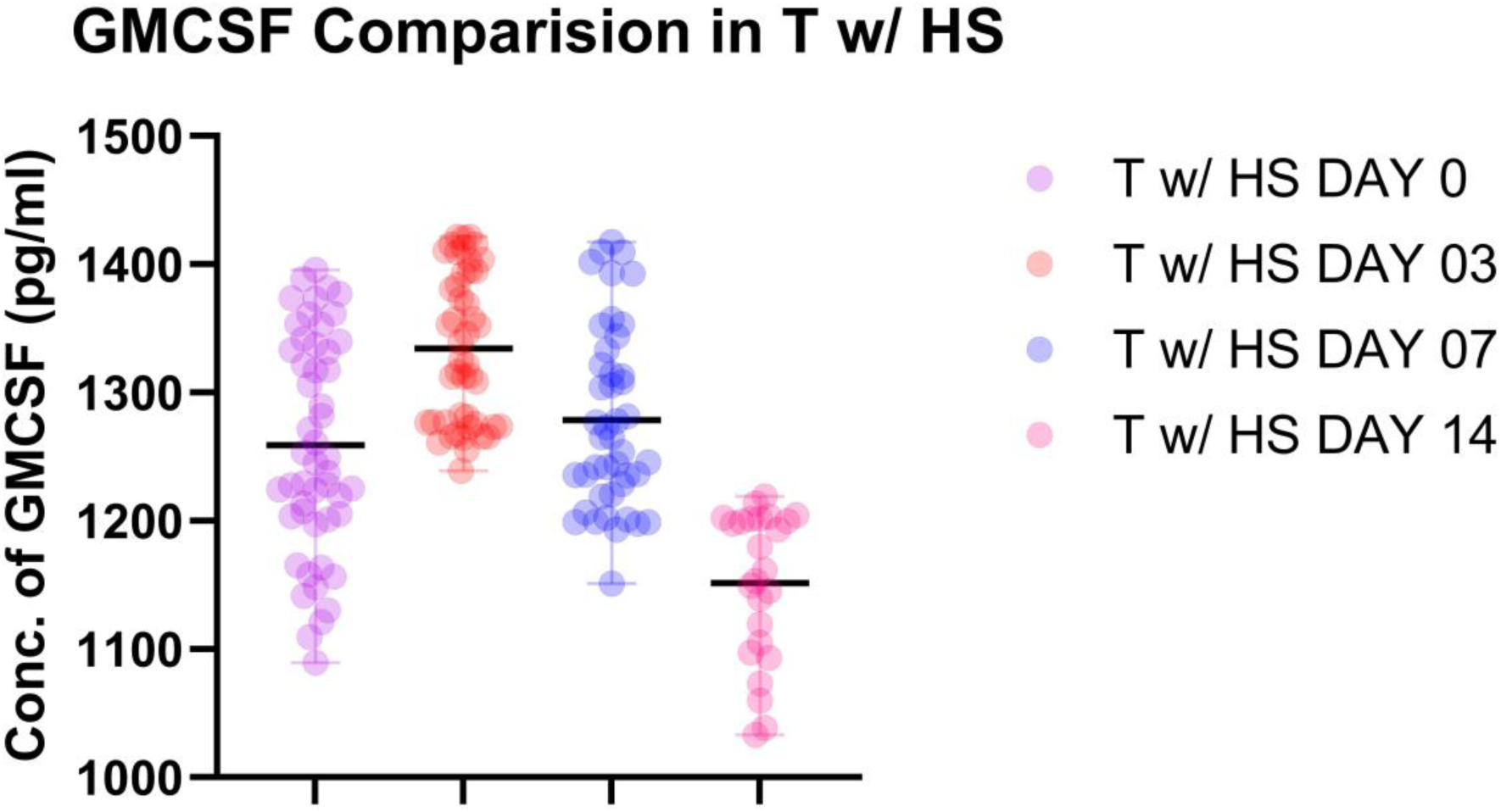
Concentration of GMCSF in the serum sample of Index Group, n=50 (Trauma Patients with Hemorrhagic Shock and Traumatic Brain Injury); ****p<0.0001.

**Figure 34:**
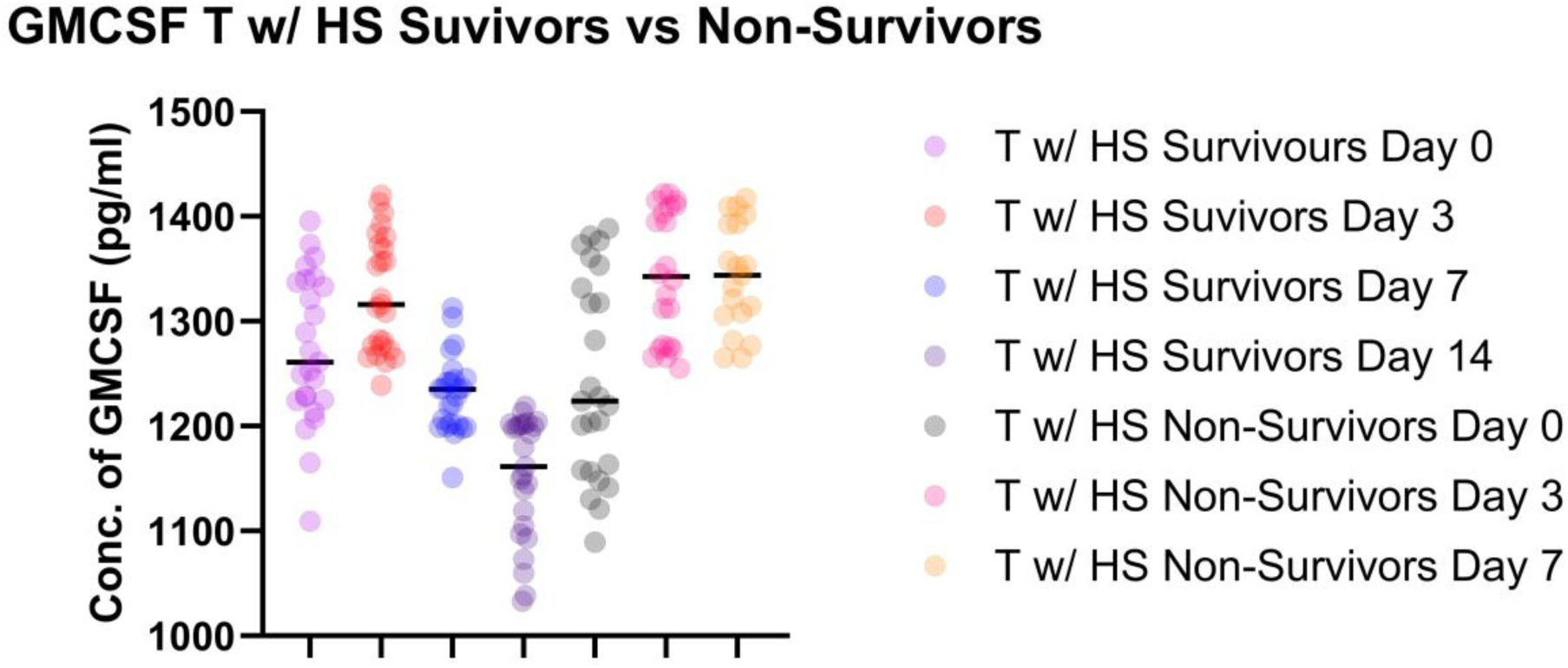
Concentration of GMCSF in serum samples among survivors and non-survivors of hemorrhagic shock, n=50 (Trauma Patients with Hemorrhagic Shock and Traumatic Brain Injury); ****p<0.0001.

Correlation analysis demonstrated that HSPC mobilization was most strongly and positively correlated with SDF-1 and VEGF-A levels, followed by G-CSF and GM-CSF, particularly during the first 72 hours post-injury (Figure 35). These cytokines are known hematopoietic chemoattractants and growth factors that mediate bone-marrow egress of progenitor cells. The correlation coefficients declined by day 7–14, aligning with the temporal reduction in circulating HSPCs. This pattern indicates that early cytokine surges likely govern acute progenitor cell mobilization, which subsequently diminishes as inflammatory stimuli subside or bone-marrow exhaustion develops.

**Figure 35:**
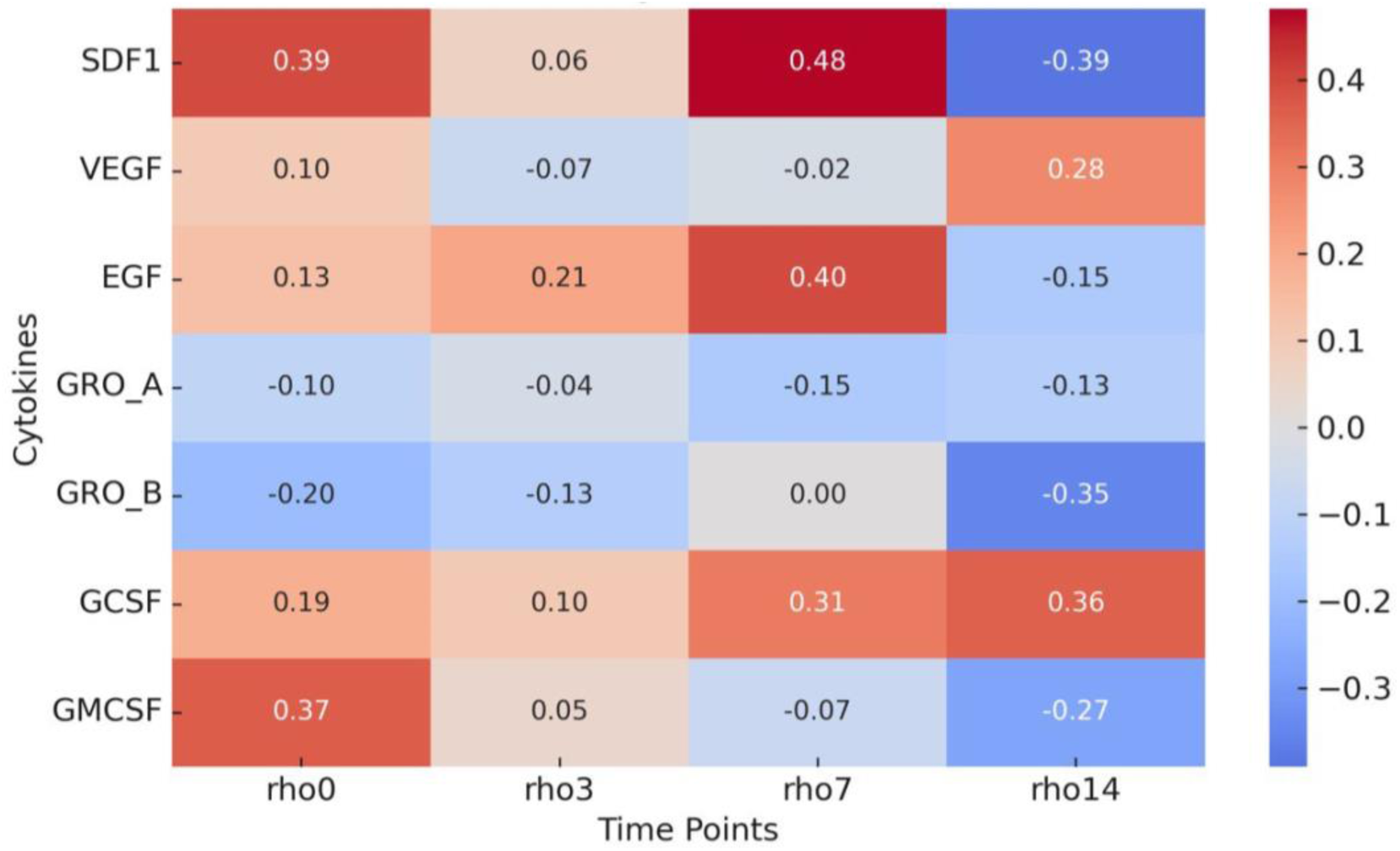
Spearman correlation heatmap between serum cytokine concentrations (SDF-1, VEGF-A, EGF, GRO-α, GRO-β, GM-CSF, and G-CSF) and hematopoietic stem/progenitor cell (HSPC) percentages across time points (day 0, 3, 7, 14) in trauma patients with hemorrhagic shock.

MSC mobilization correlated significantly with angiogenic and chemotactic cytokines; notably VEGF-A, SDF-1, and GRO-α, during the acute post-injury period, suggesting that endothelial activation and stromal recruitment are co-regulated (Figure 36). By day 7–14, these correlations weakened, consistent with resolution of the initial inflammatory and vascular signaling phase. The overall pattern implies that MSC dynamics in hemorrhagic shock are driven by early endothelial and chemokine cues rather than sustained inflammatory activity

**Figure 36:**
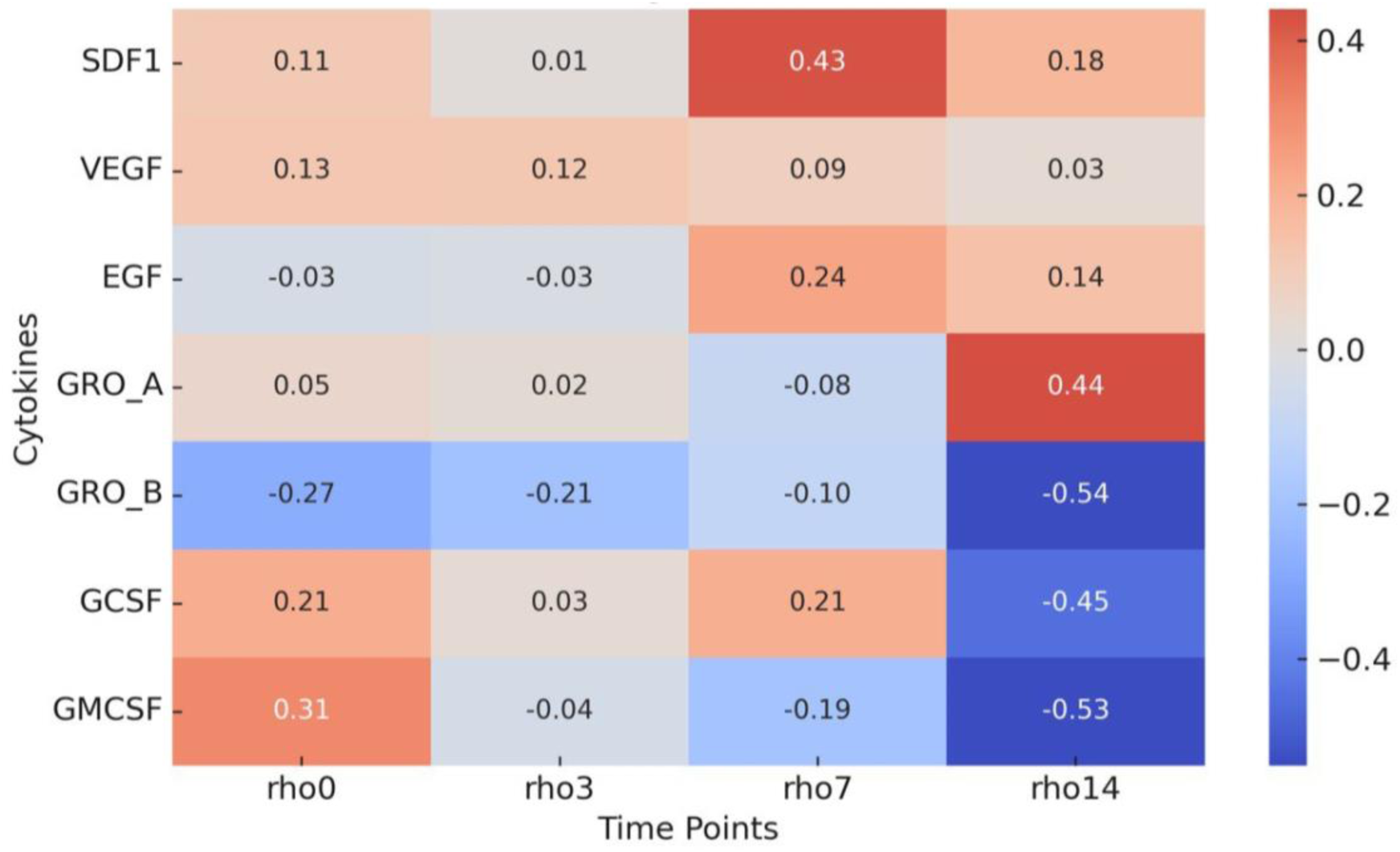
Spearman correlation heatmap between serum cytokine concentrations (SDF-1, VEGF-A, EGF, GRO-α, GRO-β, GM-CSF, and G-CSF) and mesenchymal stromal cell (MSC) percentages across time points (day 0, 3, 7, 14) in trauma patients with hemorrhagic shock.

**Figure 37:**
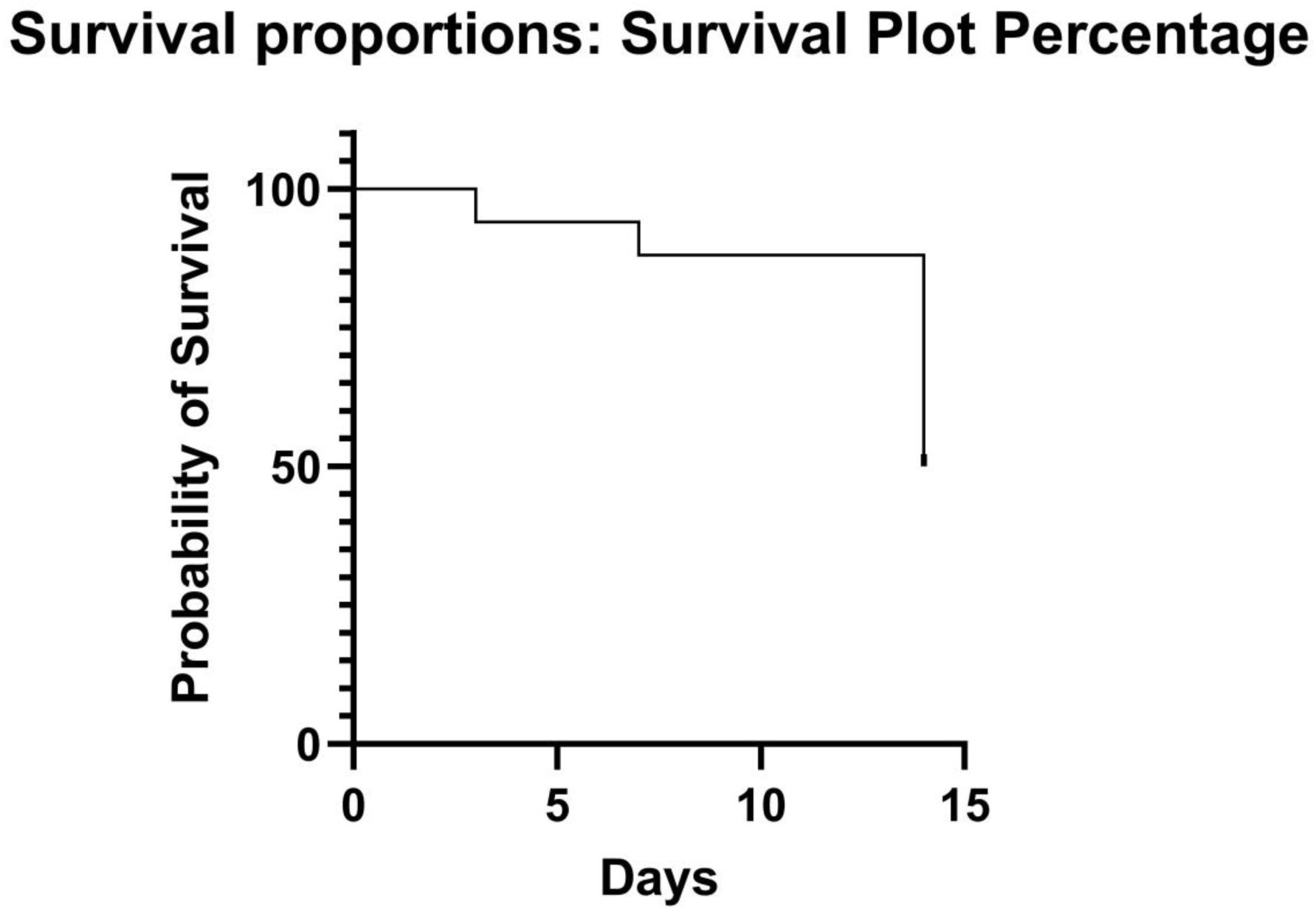
Kaplan-Meier Survival Curves of study population of Index group over 14 days, n=50 (Trauma Patients with Hemorrhagic Shock and Traumatic Brain Injury); The survival rates were 100% on Day 0, 94% at Day 03, 88% at Day 7 and 50% on Day 14.

Correlations and clinical associations

- SPC-cytokine relationships: Positive correlations were observed between key chemoattractants/hematopoietic growth factors (notably SDF-1, VEGF-A, GRO-α, G-CSF and GM-CSF) and the percentages of circulating MSC- and HSPC-like cells, consistent with a chemoattractant-driven mobilisation process.
- SPC, cytokine and organ dysfunction: Higher early SPC percentages and higher cytokine levels were associated with higher SOFA scores. Survivors typically had maximum SOFA ≤8, whereas non-survivors had progressively higher SOFA ranges (day 0–3 deaths: SOFA 10–12; day 3–7 deaths: SOFA 10–14; day 7–14 deaths: SOFA 14–22).
- SPC, cytokine and sepsis temporal association: Among non-survivors, patients who died between day 7–14 most commonly had documented late sepsis, those who died between day 3–7 showed early sepsis, and those who died between day 0–3 frequently had no conclusive signs of sepsis prior to death. Persistent elevation of SPCs and cytokines preceded or accompanied septic deterioration in the late- and early-sepsis subsets.

## Discussion

This prospective cohort study provides synchronised, serial profiling of circulating MSC- and HSPC-like populations and multiple cytokines in trauma patients with and without hemorrhagic shock over the first two weeks after injury. Three main observations emerge:

1. Trauma triggers early SPC mobilisation and cytokine up-regulation. Trauma patients without HS showed robust early increases in circulating MSCs and HSPCs and parallel cytokine elevations compared with minor-injury controls. These findings align with prior reports that tissue injury induces chemokine/cytokine signals (e.g., SDF-1α) that mobilise progenitor cells from the marrow niche (Hattori et al., 2001; Ceradini et al., 2004).
2. HS modifies SPC kinetics, early peak then decline. In HS patients, SPC mobilization was present acutely (day 0–3) but was followed by a moderate to sharp decline by days 7–14. This suggests that severe hemorrhage may transiently mobilise SPCs but either exhaust marrow reserves, impair proliferation/survival, or redistribute cells to injured tissues, leading to later depletion, consistent with prior observations of impaired HPC function after severe trauma and HS (Kumar et al., 2016; Badami et al., 2007).
3. Persistent high SPCs/cytokines associate with poor outcome. Non-survivors exhibited higher SPC percentages and cytokine levels than survivors, particularly at early time points and continuing before death. This could reflect a maladaptive or overwhelmed regenerative/immune response where high systemic inflammation and continuous mobilization are markers (or mediators) of more severe disease and impending organ failure. The observed associations with higher SOFA scores and higher rates of sepsis in non-survivors support this interpretation.

Our findings are concordant with earlier smaller studies reporting early mobilization of hematopoietic progenitors after trauma (Kumar et al., 2016) and experimental work describing MSC-mediated endothelial protection after HS (Pati et al., 2011; Potter et al., 2017). The observed correlation between chemokines (SDF-1α) and SPC mobilization supports established chemoattractant roles (Hattori et al., 2001; Wara et al., 2011). The novel contribution here is a synchronised multi-cytokine and multi-marker SPC profile across multiple time points in a larger cohort (n = 100), enabling clearer temporal patterns and outcome correlations.

Clinical and theoretical implications: The distinct temporal patterns suggest an early therapeutic window (day 0 to day 3) when exogenously administered SPCs or modulators therapies might augment endogenous repair and patients with persistent elevated SPC and cytokine signatures may be at heightened risk for deterioration and could be prioritised for intensified monitoring or targeted interventions. Additionally, SPC and cytokine profiling might provide prognostic biomarkers to stratify risk after trauma and hemorrhagic shock.

This study has limitations that should be acknowledged. The identification of stem and progenitor cells (SPCs) relied on the specific marker panel available in this study (CD123, CD38, CD45, CD105, CD73, CD90, and CD20). Although these markers effectively captured key stromal and hematopoietic subsets, complementary functional assays such as colony-forming unit analysis or lineage-tracing would provide stronger validation of true progenitor function. As this was an observational cohort study, causal relationships cannot be inferred, particularly regarding whether persistent SPC mobilization contributes to poor outcomes or merely reflects the severity of injury. In addition, potential confounders such as traumatic brain injury heterogeneity, transfusion and resuscitation practices, and infection management could have influenced SPC kinetics, despite adjustment using SOFA scoring. Finally, clinical and logistical constraints occasionally prevented sample collection at all time points, which may have introduced bias in the longitudinal trajectory estimates.

Future research should focus on elucidating the mechanistic basis of SPC mobilization in trauma and hemorrhagic shock. Specifically, studies are needed to determine whether persistent SPC mobilization directly contributes to organ dysfunction or simply reflects an inadequate repair response. Functional characterisation of mobilised cells through colony-forming assays, detailed immunophenotyping, secretome profiling, and analysis of MSC-derived extracellular vesicles would provide deeper insights into their regenerative and immunomodulatory roles. Interventional studies assessing the therapeutic potential of enhancing SPC activity or modulating key cytokines during the early post-trauma period could further clarify clinical applicability. Additionally, integrating single-cell transcriptomic and proteomic profiling would enable a more precise definition of SPC states, lineage trajectories, and their prognostic significance in trauma recovery.

## Conclusion

This prospective study demonstrates that severe trauma provokes rapid mobilisation of circulating MSC- and HSPC-like cells accompanied by broad cytokine up-regulation. In patients with hemorrhagic shock, the mobilisation response peaks within the first 72 hours and then declines by days 7–14, whereas persistent early elevation of SPCs and cytokines identifies patients at higher risk of organ dysfunction, sepsis, and death. These findings suggest that serial SPC and cytokine measurements could serve as prognostic biomarkers and highlight an early post-injury window when cell-based or cytokine-targeted therapies might be most beneficial. Further mechanistic and interventional studies, incorporating functional cell assays, expanded marker panels and single-cell profiling, are warranted to clarify causality and to translate these observations into targeted therapies for trauma care.

## Ethics Statement

This study was conducted in accordance with the Declaration of Helsinki. The study protocol was submitted for approval to the Institutional Ethics Committee of All India Institute of Medical Sciences (AIIMS), New Delhi. Ethical clearance for human subject research was applied for prior to initiation for the study. Written informed consent was obtained from all participants or their legally authorised representatives before inclusion in the study.

## Data Availability

The datasets generated and/or analysed during the current study are available from the corresponding author on reasonable request.

